# Bora, CEP192 and Cenexin activate different Plk1 pools and regulate distinct cell and centrosome cycle transitions

**DOI:** 10.1101/2025.09.30.679461

**Authors:** Devashish Dwivedi, Crisálida Borges, Daniela Harry, Luca Cirillo, Patrick Meraldi

**Author notes:** Correspondence to – and.

## Abstract

Polo-like kinase 1 (Plk1) regulates multiple steps of the cell and centrosome cycle, including mitotic entry, DNA-damage recovery, centrosome maturation and centriole disengagement. Plk1 activity depends on several independent cofactors, such as the cytoplasmic Bora, and the centrosomal proteins Cep192 and Cenexin. However, whether these Plk1 coactivators differentially regulate the Plk1-dependent processes is unknown. Here, we show that each Plk1 coactivator controls different cell cycle steps via distinct Plk1 pools in human cells. While Bora is the main driver for mitotic entry, DNA-damage recovery and centrosome maturation, centriole disengagement is mainly regulated by Cep192 and Cenexin. Moreover, we find that Plk1 and Cep192 drive S-phase progression by promoting replication origin firing. Our results thus uncover the complexity of the Plk1 activation regulatory network, in which distinct upstream activators dictate its activity in a context-dependent manner.

## Introduction

The Polo-like kinase 1 (Plk1) is a conserved serine/threonine kinase that controls key steps of the cell and centrosome cycles in animal cells to ensure faithful cell division^1^. In the mammalian cell cycle, Plk1 drives mitotic entry by phosphorylating and activating the Cdc25c1 phosphatase and inhibiting the Wee1 and Myt1 kinases^2–4^, resulting in full Cdk1-Cyclin B activation. Plk1 activity also modulates the DNA damage response, as it is essential for re-entry into the cell cycle after a DNA damage induced cell cycle arrest^5–7^. At the level of the centrosomes, Plk1 drives the dramatic increase in microtubule nucleation capacity in late G2 phase, called centrosome maturation; moreover, it is required at mitotic exit for centriole disengagement, the loss of the orthogonal arrangement of the two centrioles that allows centrosome duplication in the next cell cycle. Finally, in mitosis itself, Plk1 regulates multiple processes, which include nuclear envelope disassembly, chromosome condensation, spindle assembly and the stability of kinetochore microtubules^1, 8–10^.

Plk1 activity and abundance show a complex pattern in time and space. The protein consists of two halves, the N-terminal half with the kinase domain and the regulatory C-terminal half. This latter half contains a polo-box domain, which binds to Plk1 substrates via a phospho-epitope^11, 12^. During G1 and S-phase Plk1 shows little activity, as the C-terminal polo-box domain allosterically inhibits the kinase domain^13, 14^. Plk1 activity is mostly confined to the nucleus. Nevertheless, a centrosome-bound pool has been reported to promote DNA replication^15, 16^. At the onset of G2 phase, Plk1 activity rises sharply^2, 17^. During this phase it contributes to recovery from DNA damage, promotes mitotic entry and drives centrosome maturation. Plk1 activation requires the disruption of the autoinhibitory interaction between the kinase domain and the C-terminal half, which exposes its activation loop (T-loop) for phosphorylation at Thr-210 by the mitotic kinase Aurora-A for full activation^13, 18–20^. T-loop phosphorylation by Aurora A occurs in presence of Plk1 activators, which form a tripartite complex with Aurora A and Plk1 to provide interface for phosphorylation^6, 21^. The prototypical example is Aurora borealis/Bora^22^, which gets phosphorylated in the cytoplasm by Cdk1-Cyclin A, allowing it to bridge Aurora-A and Plk^19, 20, 23–25^. A similar bridging of Aurora-A and Plk1 is thought to occur at centrosomes via the centrosomal protein Cep192^19, 20, 26, 27^. Moreover, a second centrosomal protein, Cenexin, has also been proposed to promote Plk1 activity in G2^28–30^. Thus, different Plk1 co-activators co-exist. However, the rationale behind simultaneous utilisation of multiple activators/adaptors to activate the same kinase Plk1, remains unclear. Whether they provide redundancy as a fail-safe mechanism, fine-tune Plk1 activity in space and time, or impart functional specialisation by regulating discreet cellular Plk1 pools to control distinct cellular functions, remains unclear.

Here, we show in human cells that the Plk1 activators Bora, Cep192 and Cenexin activate different pools of Plk1 and regulate distinct functions during the cell- and centrosome-cycle progression. While all three activators contribute to the overall Plk1 activity, cytoplasmic Bora is the key driver for DNA-damage recovery, mitotic entry and centrosome maturation in late G2. In contrast, activation of Plk1 by the centrosomal protein Cep192 is required for S-phase progression and replication origin firing. Finally, Cep192- and Cenexin-dependent Plk1 activity drive centriole disengagement. These findings reveal that Plk1 activation is not governed by a singular mechanism but instead involves a network of upstream regulators that modulate its activity in a context-dependent manner.

## Results

### Different Plk1 activators regulate the activity of distinct Plk1 pools

The Polo-like kinase plays key roles during G2. During this cell cycle phase, it localizes both to the cytoplasm/nucleoplasm and is enriched at centrosomes^31^. To test the contribution of the Plk1 activators, Bora, Cep192 and Cenexin, to the activity of each pool we generated RPE1 cells expressing Fösters Resonance Energy Transfer (FRET) based Plk1 activity sensors^32, 33^ located either in the cytoplasm/nucleoplasm or at centrosomes. In addition, these cells expressed an inducible Cyclin A2-mScarlet, allowing us, based on the presence of Cyclin A2 in the cytoplasm^17^, to identify G2 cells in an asynchronous population. Applying single or combined siRNA treatments, we first found that depletion of Bora, Cep192 and Cenexin reduced Plk1 activity in the cytoplasm/nucleoplasm and at centrosomes to the same levels seen after depletion of Aurora-A (the kinase activating Plk1^22^) or treatment with a Plk1 inhibitor BI2536^34^. This confirmed that these three proteins are the main Plk1 activators (Fig. 1A-D). Second, we found that Plk1 activity in the cytoplasm/nucleoplasm was primarily, but not exclusively, dependent on Bora, whereas the centrosomal Plk1 pool was primarily dependent on the centrosomal activators Cep192 and Cenexin (Fig. 1A-D and S1A-D; Validation of depletion S1E-G). Third, immunoblotting indicated that loss of one Plk1 co-activator is not compensated by overexpression of the other ones (Fig. S1E). These results suggest that Plk1 activation by Bora, Cep192 and Cenexin occurs in a compartmentalized manner, where Bora regulates primarily the activity of the cytoplasmic/nucleoplasmic Plk1 pool and Cep192 and Cenexin regulate primarily Plk1 activity at centrosomes. Nevertheless, the fact that Plk1 activity at either pool could only be suppressed after depletion of all three co-activators, suggested continuous exchange of Plk1 between both compartments.

**Figure 1:**
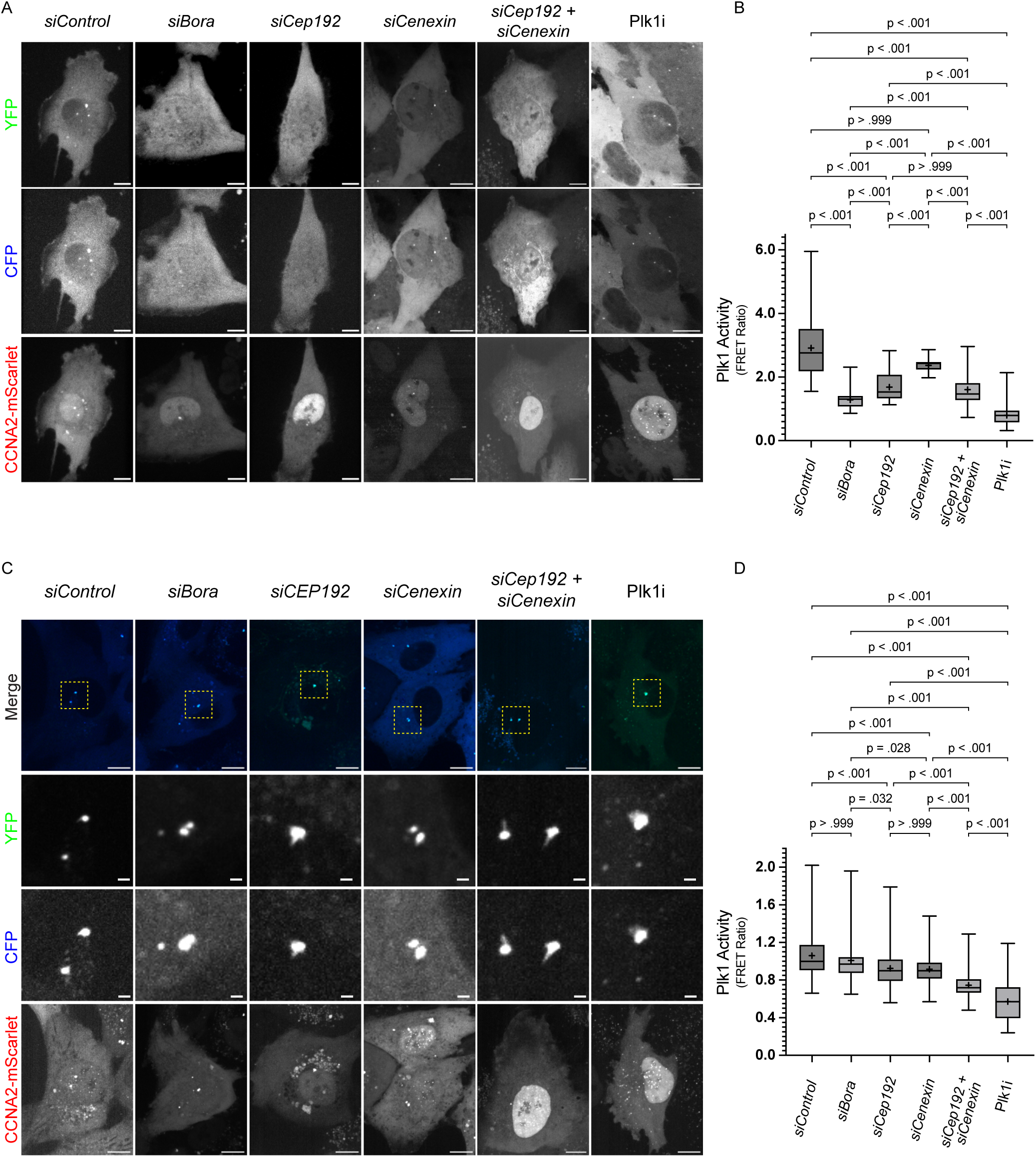
Different Plk1 activators activate distinct Plk1 pools. **(A)** Representative images of G2 phase hTERT-RPE1 cells expressing cellular Plk1 activity sensor and Cyclin A2-mScarlet treated with indicated siRNAs or inhibitors. **(B)** Box and Whisker plot for FRET measurement-based quantification of cellular Plk1 activity calculated from G2 phase cells represented in **(A)** derived from N=3 independent experiments comprising n= 115: siControl, 124: siBora, 117: siCep192, 122: siCenexin, 132: siCep192 + siCenexin and 125: Plk1i cells. **(C)** Representative images of G2 phase hTERT-RPE1 cells expressing centrosomal Plk1 activity sensor and Cyclin A2-mScarlet treated with indicated siRNAs or inhibitors. **(D)** Box and Whisker plot for FRET measurement-based quantification of centrosomal Plk1 activity calculated from G2 phase cells represented in **(C)** derived from N=4 independent experiments comprising n= 183: siControl, 183: siBora, 181: siCep192, 183: siCenexin, 185: siCep192 + siCenexin and 192: Plk1i cells. Data presented as the entire range with values between first and third quartiles in boxes. ‘**+**’ sign in each column represent the mean value (p values for individual comparisons indicated on graphs: Kruskal-Wallis test). Scale bars = 5 µm.

### Plk1 localisation is dynamic on the centrosome

To test whether Plk1 exchanges between the cytoplasmic/nucleoplasmic pool and the centrosomal pool, we recorded its recovery dynamics on centrosomes after photobleaching (FRAP) in RPE1 cells expressing endogenously tagged GFP-Plk1. FRAP-recordings of control-depleted cells in which centrosomal GFP-Plk1 was bleached, indicated the presence of at least two GFP-Plk1 pools: 2/3 of GFP-Plk1 rapidly exchanged with the cytoplasmic pool with (t_1/2_ < 10s), while 1/3 of GFP-Plk1 was immobile over a period of 3 minutes (Fig. 2A-B and Supplementary Video 1). To test the contribution of Cep192 and Cenexin to this immobile pool, we depleted either protein or both in combination. The immobile fraction diminished upon depletion of either centrosomal co-activator and disappeared after Cep192/Cenexin co-depletion (Fig. 2C-I and Supplementary Videos 2-4). This change in the mobility was due to a change in Plk1 anchoring rather than Plk1 activity itself, since Plk1 inhibition had no effect on the relative proportion of mobile and immobile Plk1 (Fig. 2J and S2A-D and Supplementary Video 5-6). Finally, we also tested whether Bora, Cep192 or Cenexin-depletion affected the overall abundance of GFP-Plk1 at centrosomes and found that all three depletions caused a moderate reduction in Plk1 levels (Fig. 2K-L and S2E-F), consistent with the previous observation that Plk1 activity can regulate its centrosomal abundance. We conclude that, in G2, the majority of the centrosomal pool of Plk1 exchanges with the cytoplasm/nucleoplasm, while a smaller immobile fraction likely binds to Cep192 and Cenexin.

**Figure 2:**
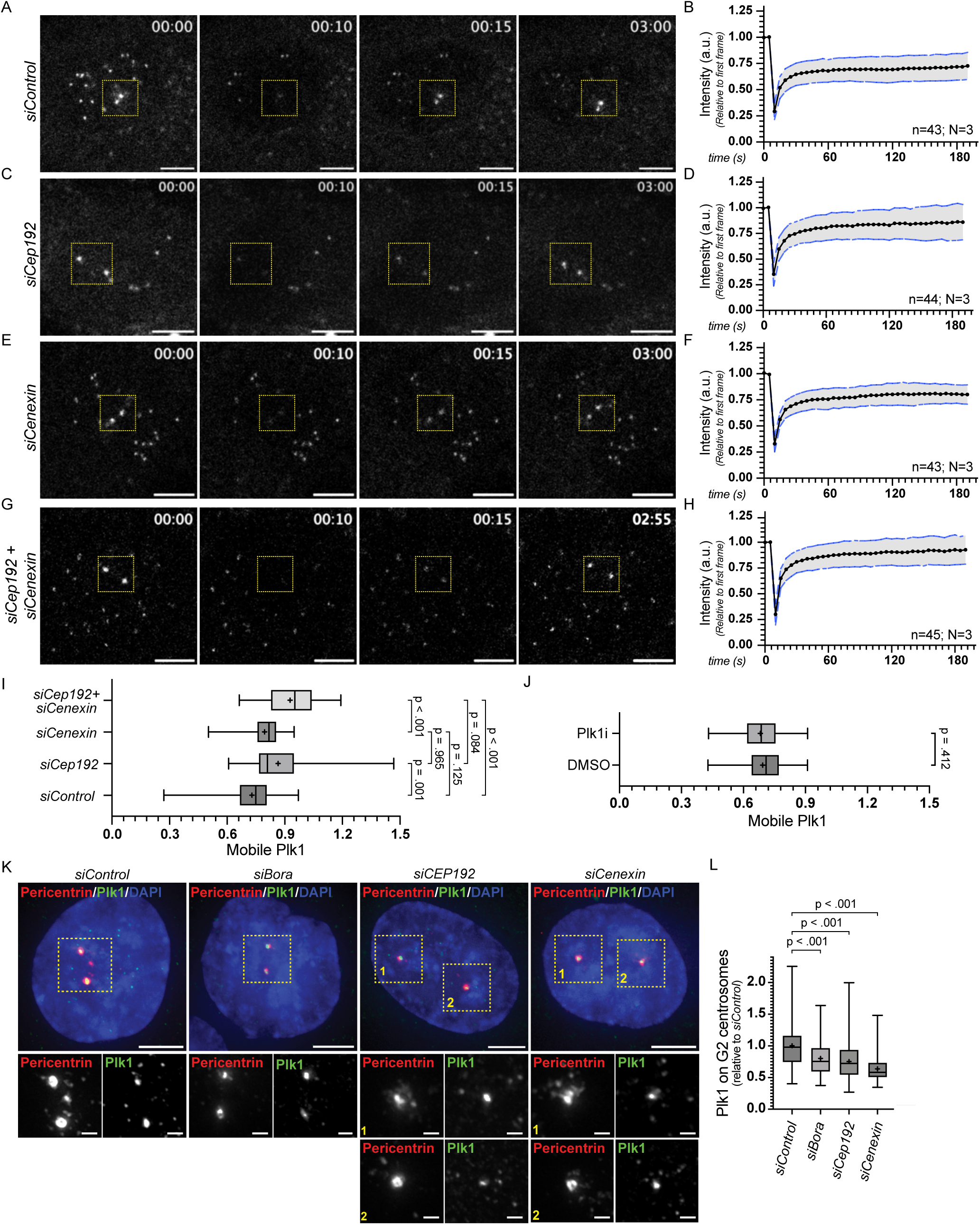
Plk1 dynamically localises on centrosomes. **(A-H)** Representative time stamp images and quantifications from FRAP movies from G2 phase Plk1-EGFP tagged hTERT-RPE1 cells treated with non-targeting siRNA control **(A;** quantification of recovery in **(B))**, Cep192 siRNA **(C** quantification of recovery in **(D))**, Cenexin siRNA **(E** quantification of recovery in **(F))** and siCep192 + siCenexin **(G** quantification of recovery in **(H))**. **(I)** Box and Whisker plot for measurement of mobile Plk1 pool on centrosomes under indicated conditions (N=3 independent experiments; n= 43, 44, 43 and 45 for siControl, siCep192, siCenexin and siCep192 + siCenexin, respectively.) **(J)** Box and Whisker plot for measurement of mobile Plk1 pool on centrosomes after Plk1 inhibition derived from N=3 independent experiments comprising n= 43, and 53 individual cells for DMSO and Plk1i, respectively. **(K)** Representative immunofluorescence images of hTERT-RPE1: Plk1-EGFP cells treated with indicated siRNAs and stained for Pericentrin (Red) and EGFP (Green). **(L)** Box and Whisker plot for measurement of Plk1 levels on centrosomes during G2 phase under the conditions indicated in **(K)** generated from N=5 independent experiments comprising n= 287: siControl, 279: siBora, 276: siCep192 and 287: siCenexin cells. Data presented as the entire range with values between first and third quartiles in boxes. ‘**+**’ sign in each column represent the mean value (p values for individual comparisons indicated on graphs: Kruskal-Wallis test). Scale bars Images: 5 µm and zoomed insets: 0.5 µm.

### Plk1 activity on the centrosomes is essential for timely progression of S-phase

Plk1 activity is crucial for progression through different cell cycle stages^1^. We therefore investigated how the depletion of Bora, Cep192 and Cenexin affects the duration of S and G2 phases. We performed long-term time lapse imaging of live RPE1 cells expressing endogenously tagged mRuby-PCNA, which enabled us to distinguish between the different cell-cycle phases. Control-depleted cells completed S phase in 3.88±1.21 h and G2 in 4.59±1.32 h (Fig. 3A-B and 4A-B; Supplementary Video 7 and 8). Depletion of Cyclin A2, a critical regulator of S-and G-phase and our positive control, strongly delayed S-phase and almost completely blocked the G2/M transition, as 95% of the cells failed to enter mitosis (Fig. 3A-B, and 4A-B; Supplementary Video 9 and 10). Depletion of any of the three Plk1 co-activator significantly increased S-phase duration, with Cep192 depletion causing the strongest delay (9.57±2.85 h; Fig. 3A-B; Supplementary Video 11-13). In contrast only Bora depletion significantly delayed G2 completion (13.99±6.48 h; Fig. 4A-B, Supplementary Video 19-21). Co-depletion of the other co-activators did not further delay S-phase compared to Cep192 depletion, nor did it further extend G2 completion compared with Bora depletion (Fig. S3 A-D; Supplementary Video 14-26). Visual inspection of the mRuby-PCNA signal, which reflects replication origin firing in S-phase^35^, suggested a reduction in this activity. Moreover, previous studies indicated that *Xenopus laevis* Plk1 regulates DNA replication-origin firing in embryonic extracts^36–38^. To test if this is also the case in human somatic cells, we depleted Bora, Cep192 or Cenexin, synchronised the cells at G1/S boundary with the Cdk4/6 inhibitor Palbociclib (Cdk4/6i)^39^, released them for 3 hrs to reach mid S phase, stained for PCNA, and counted the number of foci per nucleus. While Cep192 depletion caused roughly a four-fold reduction in PCNA foci, Bora depletion had a mild effect and Cenexin depletion had none (Fig 3C and D). Overall, we conclude that the cytosolic Plk1 activator Bora is rate-limiting for the timely progression of G2 phase, whereas the centrosomal Plk1 activator Cep192 is critical for replication origin firing and timely completion of S-phase.

**Figure 3:**
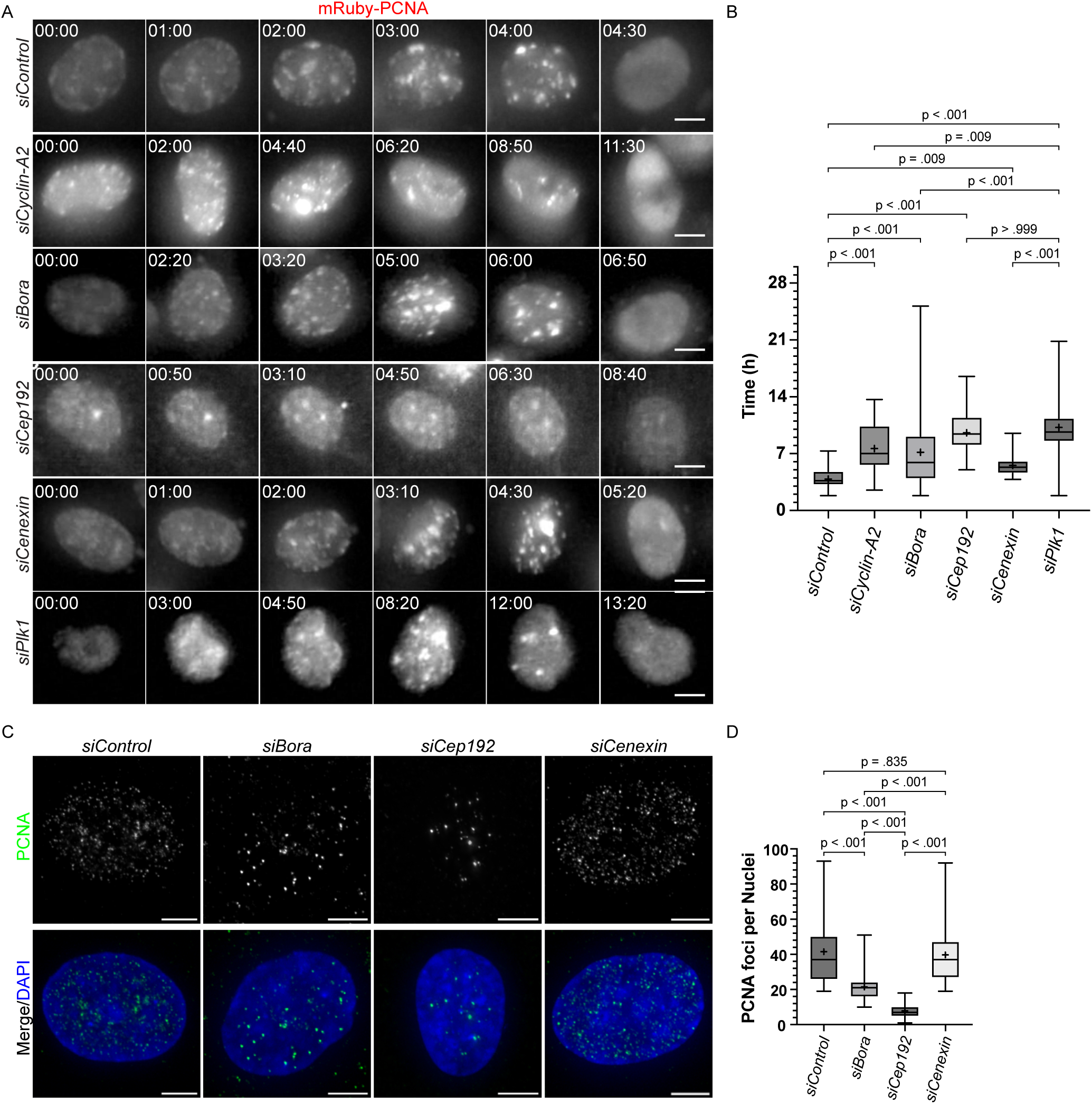
Depletion of Cep192 delays S phase progression. **(A)** Live-cell time-lapse images of hTERT-RPE1: mRuby-PCNA cells treated with indicated siRNAs as they progress through S phase. **(B)** Box and whisker plot for measurement of the duration of S phase in cells after depletion of proteins indicated in **(A)**. Data derived from N=3 independent experiments comprising n= 71: siControl, 52: siCyclin A2, 68: siBora, 46: siCep192, 57: siCenexin and 62: siPlk1 cells. **(C)** Representative images of synchronised S phase RPE1 cells treated with indicated siRNAs and immunolabelled for replication forks using PCNA antibody. **(D)** Box and Whisker plot for measuring number of PCNA foci per cells as proxy for the rate of DNA replication in conditions indicated in **(C)** derived from N= 3 independent experiments comprising n= 123: siControl, 122: siBora, 126: siCep192 and 127: siCenexin cells. Data represent the entire range with values between first and third quartiles in boxes. ‘**+**’ sign in each column represent the mean value (p values for individual comparisons indicated on graphs: Kruskal-Wallis test). Scale bars: 5 µm.

**Figure 4:**
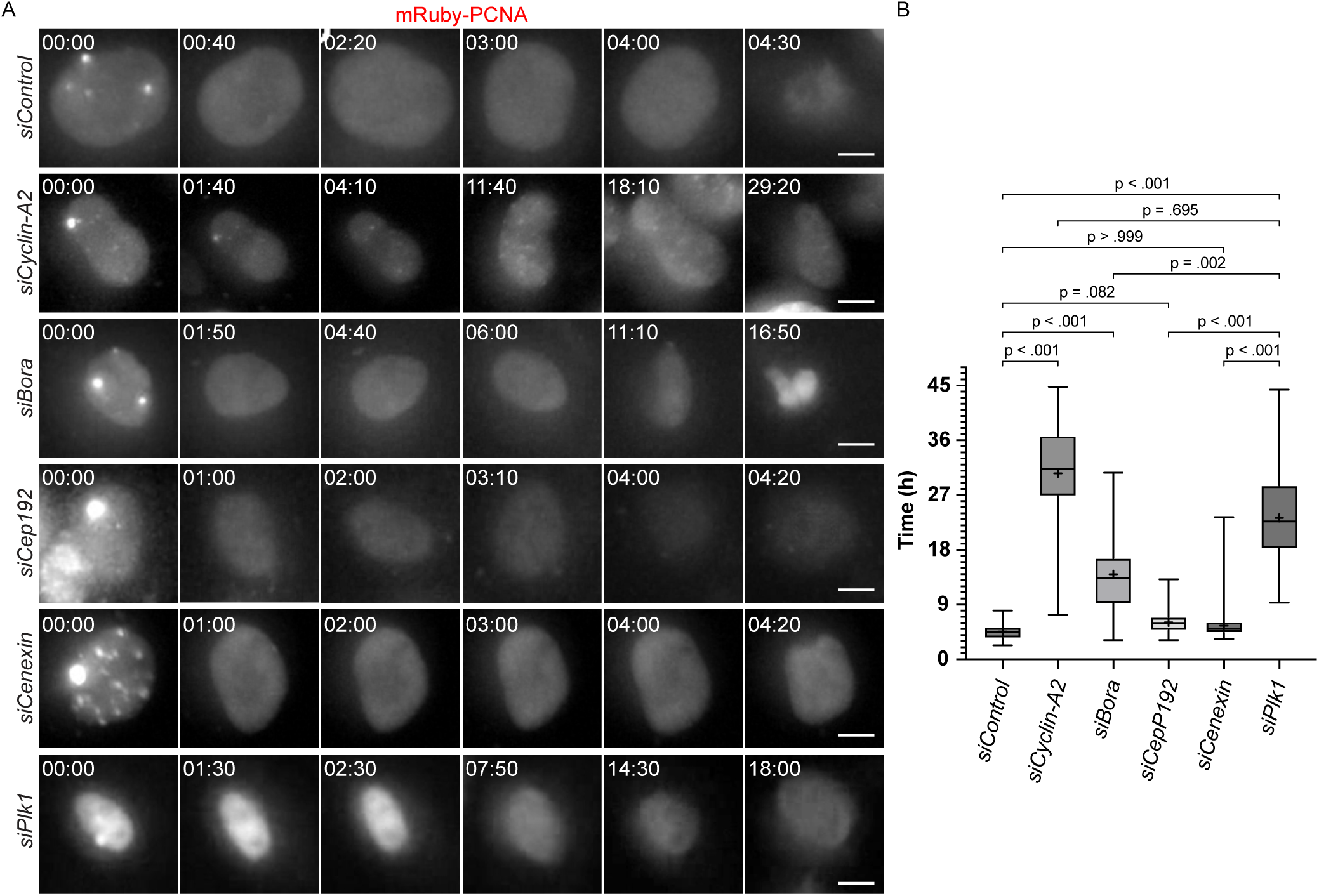
Bora depletion imposes G2 delay. **(A)** Live-cell time-lapse images of hTERT-RPE1: mRuby-PCNA cells treated with indicated siRNAs as they progress through G2 phase. **(B)** Box and whisker plot for measurement of the duration of G2 phase in cells after depletion of proteins indicated in **(A)**. Data derived from N=3 independent experiments comprising n= 83: siControl, 69: siCyclin A2, 69: siBora, 65: siCep192, 83: siCenexin and 70: siPlk1 cells. Data presented as the entire range with values between first and third quartiles in boxes. ‘**+**’ sign in each column represent the mean value (p values for individual comparisons indicated on graphs: Kruskal-Wallis test). Scale bars: 5 µm.

### Cep192 regulates both cellular and centrosomal Plk1 activity during S phase

Cep192 acts as a scaffold for Aurora A-dependent Plk1 activation^19, 20, 26, 27^, but has also been shown to affect centrosome duplication during S-phase^40, 41^. To distinguish if the prolonged S phase observed after Cep192 depletion is due to centriole duplication defects or due to loss of Plk1 activity, we generated cells without centrosomes. Specifically, we blocked centrosome duplication with the Plk4 inhibitor centrinone^42^ in RPE1 cells lacking USP28, a component of the mitotic stopwatch checkpoint that induces a p53-dependent cell cycle arrest after centrosome loss^43^. After one week of centrinone treatment and a single cell selection, we obtained a RPE1 *USP28^-/-^* cell line devoid of centrosomes (Fig. S4A and B). Flow cytometry indicated the same G1-, S-and G2/M proportions in RPE1, RPE1 *USP28^-/-^* and RPE1 *USP28^-/-^* cells lacking centrosomes (Fig. 5A-C). Nevertheless, Cep192 depletion in RPE1 *USP28^-/-^* lacking centrosomes still led to an S-phase enrichment (Fig. 5D), suggesting that Cep192 drives the S-phase progression via Plk1 activity and independently of centrosomes.

**Figure 5:**
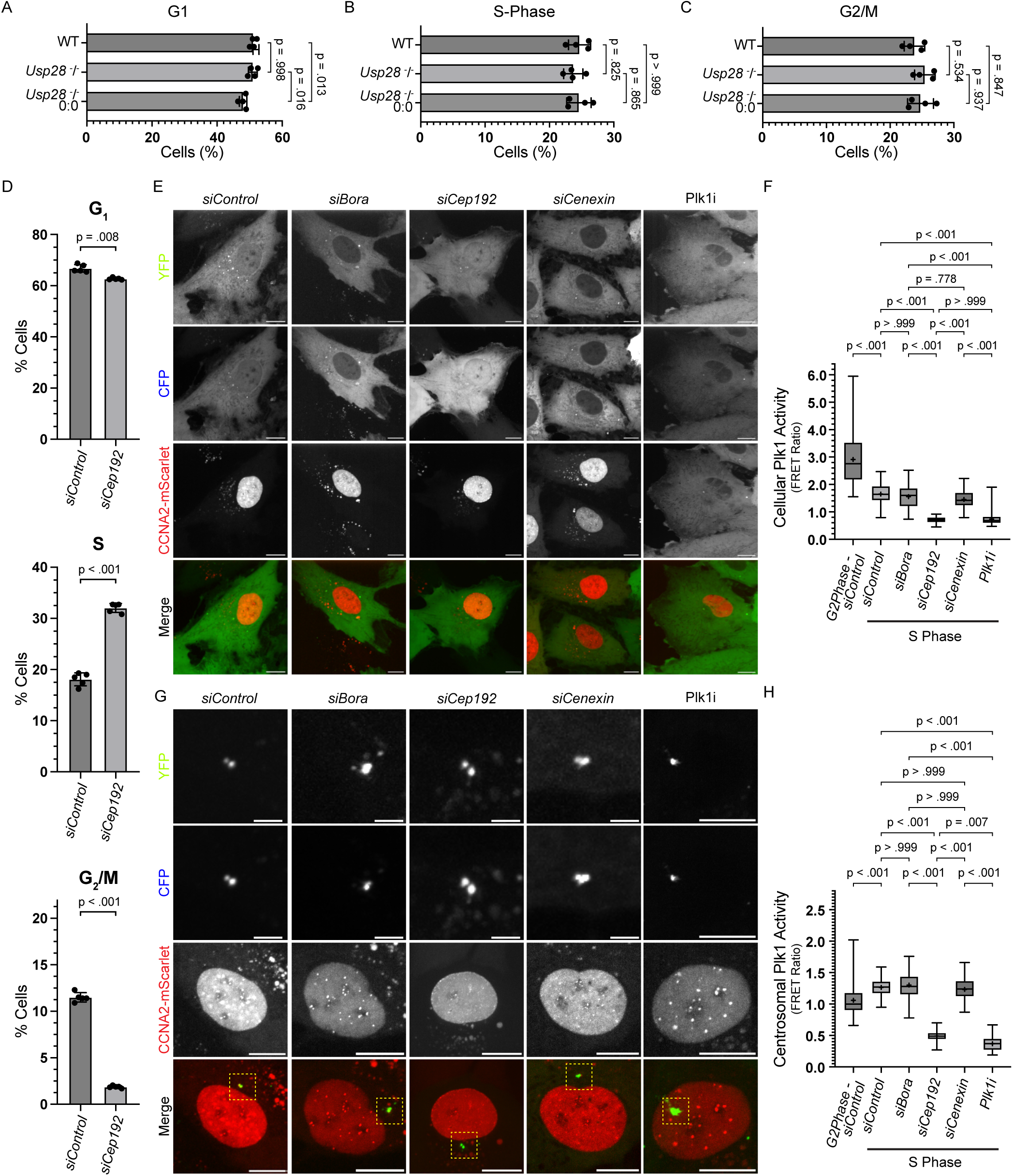
Cep192 but not Bora or Cenexin regulates Plk1 activation during S phase. **(A-C)** Flow cytometry-based quantification of cell cycle profile of hTERT-RPE1 (WT), hTERT-RPE1: Usp28 ^-^/^-^ and hTERT-RPE1: Usp28 ^-^/^-^ 0:0 cells revealing percentage of cells in G1 **(A)**, S **(B)** and G2 **(C)** phases derived from N=4 independent experiments with each treatment comprising around 30,000 cells. **(D)** Flow cytometry-based quantification of cell cycle profile or RPE1 cells treated with Control vs Cep192 siRNA showing percentage of cells in G1, S and G2 phases obtained from N=5 independent experiments with each treatment comprising of 30,000 cells. **(E)** Representative images of S phase hTERT-RPE1 cells expressing cellular Plk1 activity sensor and Cyclin A2-mScarlet treated with indicated siRNAs or inhibitors. **(F)** Box and Whisker plot for FRET measurement-based quantification of cellular Plk1 activity calculated from S phase cells represented in **(E)** derived from N=4 independent experiments comprising n= 203: siControl, 204: siBora, 215: siCep192, 219: siCenexin, 215: Plk1i cells. The leftmost **G2 Phase – siControl** bar shown is replica of siControl data shown in Fig 1B for side-by-side comparison purposes. **G)** Representative images of S phase hTERT-RPE1 cells expressing centrosomal Plk1 activity sensor and Cyclin A2-mScarlet treated with indicated siRNAs or inhibitors. **(H)** Box and Whisker plot for FRET measurement-based quantification of cellular Plk1 activity calculated from S phase cells represented in **(G)** derived from N=3 independent experiments comprising n= 163: siControl, 187: siBora, 187: siCep192, 184: siCenexin, 178: Plk1i cells. The leftmost **G2 Phase – siControl** bar shown is replica of siControl data shown in Fig 1B for side-by-side comparison purposes. Data presented as the entire range with values between first and third quartiles in boxes. ‘**+**’ sign in each column represent the mean value (p values for individual comparisons indicated on graphs: Kruskal-Wallis test). Scale bars, Images: 5 µm and zoomed insets: 0.5 µm.

We therefore quantified the cellular and centrosomal Plk1 activity during S phase after Bora, Cep192 or Cenexin depletion. Using the nuclear localisation of Cyclin A2 as proxy^17^ to identify S phase cells, we first found that the cytoplasmic Plk1 activity was much lower in S-phase compared to G2, while the centrosomal Plk1 activity was equivalent (Figure 5E-H). Moreover, Plk1 activity in S-phase at either site depended on Cep192, but not on Bora or Cenexin (Figure 5E-H). This indicated that Cep192 is the principal activator of both cytoplasmic and centrosomal Plk1 during S-phase (Figure 5E-H).

### Bora is essential for DNA-damage recovery

In addition to its generic cell cycle roles, Plk1 is also critical for cell cycle re-entry after DNA damage, as it directly inhibits the DNA-damage checkpoint effectors 53BP1, p53 and their downstream checkpoint kinases Chk1 and Chk2^44–47^. To determine which Plk1 activator regulates checkpoint recovery after DNA damage, we depleted Bora, Cep192 and Cenexin, synchronised the cells at the G1/S boundary with a Cdk4/6i, released them for 6 hours in the presence or absence of a short pulse of the DNA-damaging agent Doxorubicin, and treated them with the Eg5 inhibitor STLC for 16 hours to arrest them in mitosis (See scheme Fig. 6A). By determining the mitotic index, we could monitor the ability of cells to recover from a DNA-damage induced cell cycle arrest. Consistent with previous studies, Bora depletion specifically abolished mitotic entry after a Doxorubicin pulse, indicating a failure to recover from a DNA-damage-induced arrest (Fig. 6B-C; Confirmation of DNA damage in Fig. S6A). In contrast, Cenexin depletion did not prevent mitotic entry (Fig. 6B-C). Cep192 depletion and complete Plk1 inhibition gave inconclusive results, as these perturbations already prevented mitotic entry in the absence of Doxorubicin pulse (Fig. 6D-E and S6B-C), likely reflecting their critical contribution to S-phase progression. We therefore conclude that Bora-dependent Plk1 activity, but not Cenexin, is required for DNA-damage recovery.

**Figure 6:**
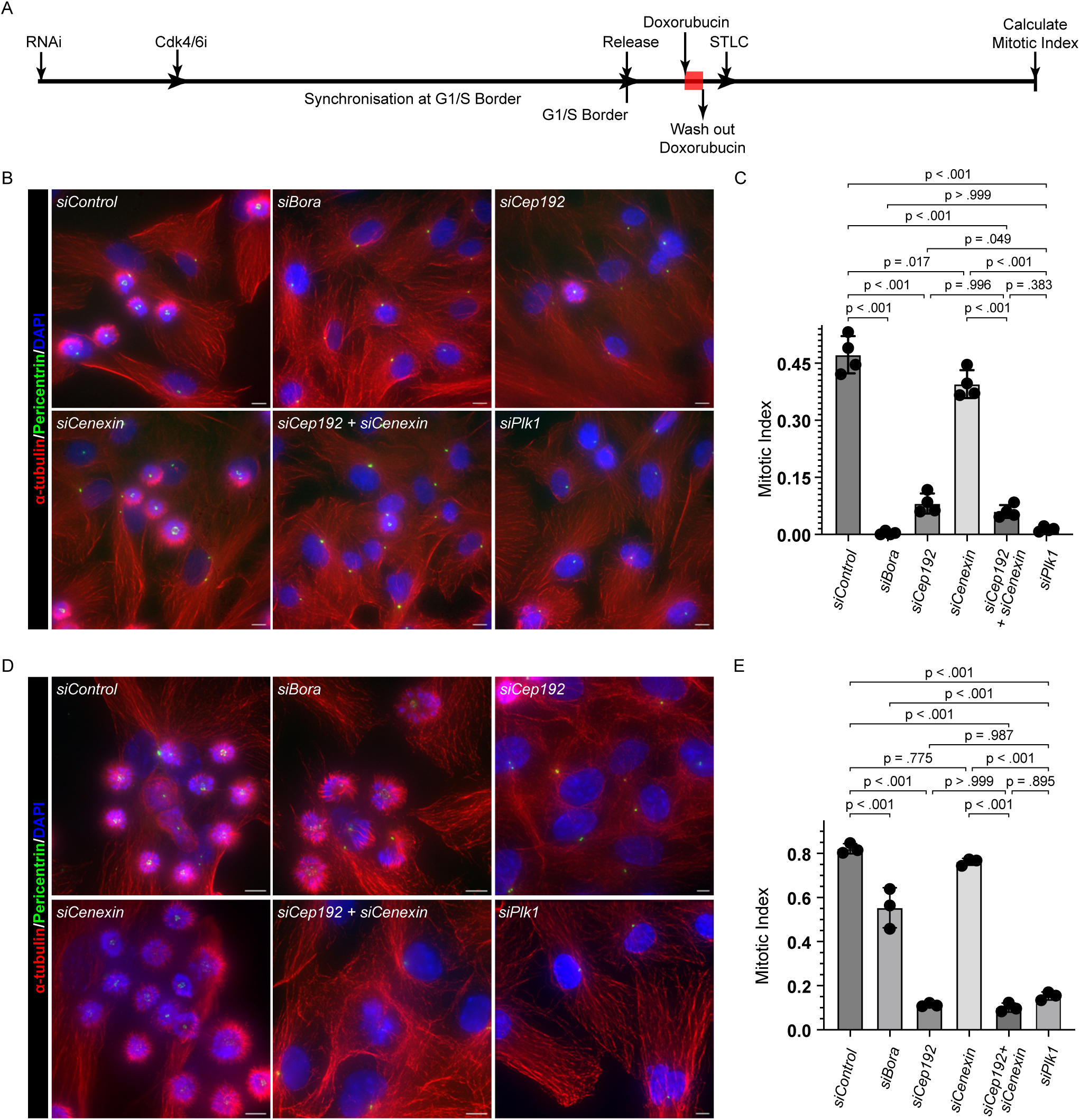
Bora but not Cep192 or Cenexin dependent Plk1 activation during G2 is required for checkpoint recovery. **(A)** Schematic representation of experimental setup for investigating the role of Plk1 activators in checkpoint recovery. **(B)** Representative field images of hTERT-RPE1 cells treated with indicated siRNAs and stained for α-tubulin, Pericentrin and DAPI. **(C)** Quantification representing mitotic index after DNA damage recovery in cells under depletion different conditions shown in **(B)** obtained from N=4 independent experiments consisting of n=1136: siControl, 1398: siBora, 976: siCep192, 1359: siCenexin, 1157: siCep192 + siCenexin and 972: siPlk1 cells. **(D)** Representative field images of hTERT-RPE1 cells treated with indicated siRNAs and stained for α-tubulin, Pericentrin and DAPI. **(E)** Quantification representing mitotic index in cells without doxorubicin pulse under different depletion conditions shown in **(D)** obtained from N=3 independent experiments consisting of n=1413: siControl, 838: siBora, 827: siCep192, 1412: siCenexin, 636: siCep192 + siCenexin and 1803: siPlk1 cells. Data presented as the entire range with values between first and third quartiles in boxes. ‘**+**’ sign in each column represent the mean value (p values for individual comparisons indicated on graphs: Kruskal-Wallis test). Scale bars: 5 µm.

### Bora is the major driver of Plk1 dependent centrosome maturation during late G2

As cells prepare for mitotic entry in late G2, Plk1 drives centrosome maturation via the recruitment of pericentrosomal matrix (PCM) proteins such as Pericentrin (PCNT) and γ-tubulin, to increase the microtubule-nucleation capacity of the future spindle poles^48–51^. To identify the contribution of each of the Plk1-adaptors in this process, we monitored the total volume of γ-tubulin and PCNT on the early prometaphase spindle poles after Bora, Cep192 or Cenexin depletion. Whereas Bora-depletion reduced the levels of γ-tubulin and PCNT on spindle poles as severely as Plk1 depletion, Cep192 depletion only caused an intermediate (50%) reduction, and Cenexin depletion had no effect (Fig. 7A-C and S7A-C). These data indicated that the cytoplasmic pool of Plk1 activated by Bora, is the key regulator of centrosome maturation.

**Figure 7:**
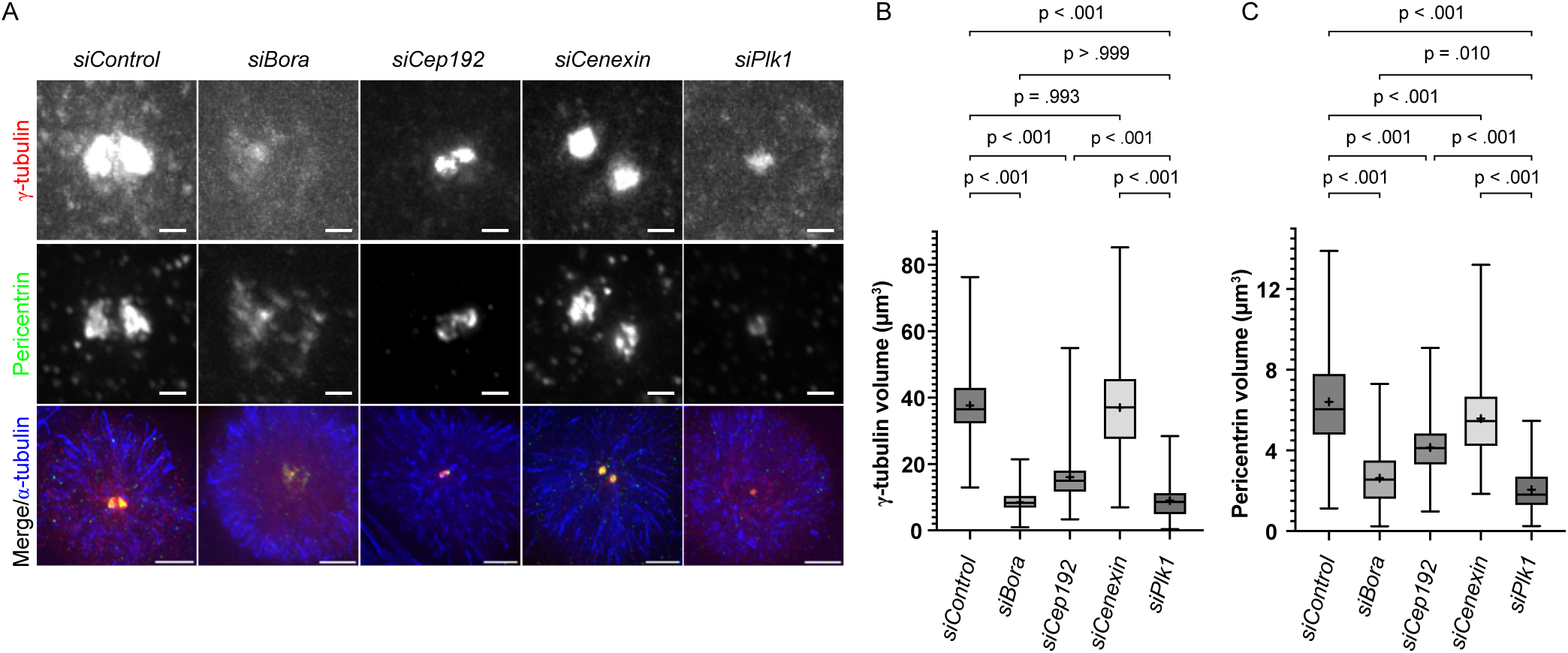
Bora and Cep192 regulate centrosome maturation. **(A)** Representative images of G2 phase centrosomes stained with γ-tubulin (red) and Pericentrin (green) in hTERT-RPE1 cells treated with indicated siRNAs. **(B-C)** Box and Whisker plots representing the centrosomal levels of γ-tubulin **(B)** and Pericentrin **(C)** in cells represented in **(A)** as derived from N=3 independent experiments consisting of n= (168: siControl, 171: siBora, 177: siCep192, 165: siCenexin and 168: siPlk1 for γ-tubulin) and (n= 176: siControl, 170: siBora, 179: siCep192, 157: siCenexin and 183: siPlk1 for Pericentrin (green) cells. Data presented as the entire range with values between the first and third quartiles in boxes. ‘**+**’ sign in each column represent the mean value (p values for individual comparisons indicated on graphs: Kruskal-Wallis test). Scale bars: 0.5 µm.

### Cep192 and Cenexin drive centriole disengagement

In cells exiting mitosis, Plk1 regulates the first step of centrosome duplication by inducing centriole disengagement during mitotic exit^52–54^. In our previous work, we showed that incomplete Plk1 activation caused by mild DNA replication stress leads to premature centriole disengagement in G2^55^. We hypothesized that such partial Plk1 activity mimics the condition found during mitotic exit, when Plk1 activity declines^56^. To analyse how this process is controlled, we depleted each of three Plk1 activator, and examined the centriole engagement status in G2 phase using Ultrastructure-Expansion Microscopy (U-ExM), as previously described ^55^.

Strikingly, Bora depletion led to premature centriole disengagement in nearly all cells (98.10±1.65% vs. 1.76±1.61% in control-depleted cells), while Cep192 depletion produced an intermediate effect (53.10±3.95% of cells with disengaged centrioles), and Cenexin loss did not induce centriole disengagement (Fig. 8A-C). Premature centriole disengagement was not due to DNA replication stress, since, unlike the DNA-polymerase inhibitor aphidicolin, none of the depletions increased typical DNA replication stress markers, such as 53BP1 or γ-H2AX foci (Fig. S8A-C). These results suggested that in conditions with partial cytoplasmic Plk1 activity but high centrosomal Plk1 activity, as seen after Bora depletion, centrioles disengage (Fig 1). To test this hypothesis, we co-depleted Bora and CEP192 either in WT RPE1 cells or in RPE1 *CENEXIN^-/-^* knock-out cells (to avoid applying a technically challenging triple siRNA treatment). Loss of either centrosomal Plk1 activator reduced centriole disengagement in a Bora siRNA background, and depletion of both almost completely abolished it (Fig. 8A-C). We conclude that centriole disengagement is primarily driven by the centrosomal Plk1 activators Cep192 and Cenexin, and that a condition with partial cytoplasmic but high centrosomal Plk1 activity in G2, is sufficient to trigger premature centriole dis-engagement.

**Figure 8:**
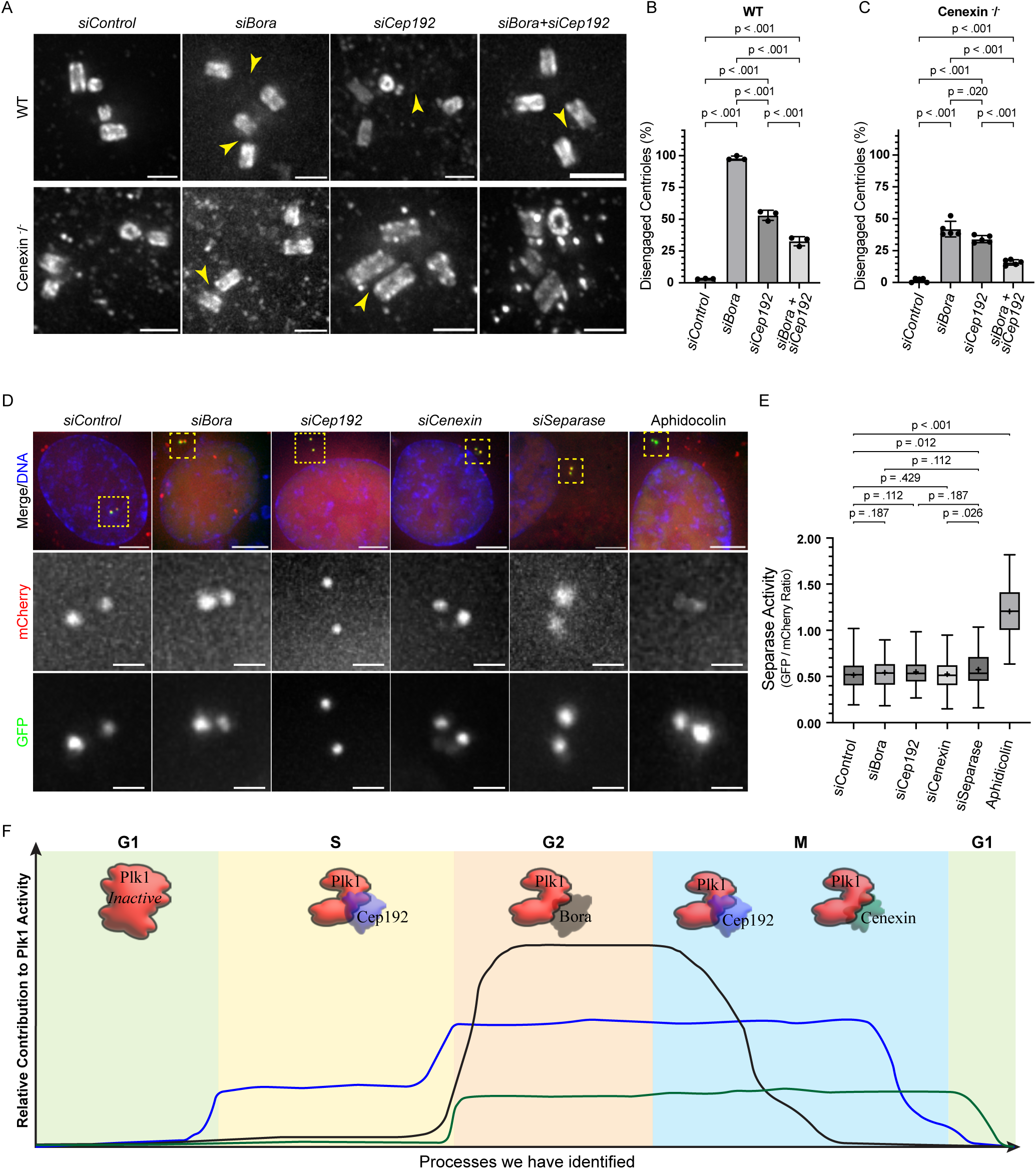
Cenexin dependent Plk1 activation drives centriole disengagement. **(A)** Expansion microscopy images of centrioles in G2 phase hTERT-RPE1 (WT) and hTERT-RPE1 Cenexin ^-^/^-^ cells stained against α-tubulin and treated with indicated siRNAs to visualise centriole configuration. The arrowheads indicate the disengaged mother daughter centriole pairs. **(B)** Quantification of percentage of G2 phase hTERT-RPE1 cells with disengaged centrioles in their centrosomes (*N* = 3 independent experiments, n = 96, 98, 100 and 104 cells in siControl, siBora, siCep192 and siBora + siCep192, respectively) **(C)** Quantification of percentage of G2 phase hTERT-RPE1 Cenexin ^-^/^-^ cells with disengaged centrioles in their centrosomes (*N* = 5 independent experiments, n = 166, 149, 138 and 145 cells in siControl, siBora, siCep192 and siBora + siCep192, respectively) **(D)** Representative images of hTERT-RPE1: Centrosomal Separase activity sensor cells depicting centrosomal intensity of GFP and mCherry after indicated siRNA/drug treatments. **(E)** Box and Whisker plot for quantification of centrosomal activity of Separase in conditions depicted in **(D)** and derived from N=3 independent experiments consisting of n=142: siControl, 144: siBora, 134: siCep192, 152: siCenexin, 152: siSeparase and 147: Aphidicolin treated cells. Data presented as the entire range with values between first and third quartiles in boxes. ‘**+**’ sign in each column represent the mean value (p values for individual comparisons indicated on graphs: Kruskal-Wallis test). Scale bars: 0.5 µm. **(F)** Schematic representation of regulation of Plk1 activity by Cep192, Bora and Cenexin during S, G2 and M phase, respectively.

Finally, we quantified the proteolytic Separase activity in the depletion leading to premature centriole disengagement. Indeed, together with Plk1, Separase plays a key role in centriole disengagement during telophase^57–59^ and in G2 following mild replication stress^55^. Using an established Separase activity sensor^60^, we monitored Separase activity on centrosomes during G2 phase, and found that, unlike mild replication stress induced by low doses of aphidicolin (positive control), none of the tested conditions upregulated Separase activity (Fig. 8D-E). Thus, centriole disengagement observed after Bora and Cep192 depletion solely rely on partial Plk1 activity.

## Discussion

Here we investigated how the different Plk1 co-activators regulate Plk1 activity in time and space and whether they differentially control the Plk1-dependent transitions of the cell-and centrosome-cycle. In terms of spatial regulation, our results indicate that the centrosomal and cytoplasmic Plk1 pools are under the control of distinct co-activators, but that a partial exchange exists between the two populations. At the functional level, we demonstrate that the Plk1-dependent cell-and centrosome-cycle steps are differentially regulated by each Plk1 co-activator. Cytoplasmic Bora is the main driver for mitotic entry, DNA-damage recovery and centrosome maturation, while centriole disengagement is controlled by the centrosomal co-activators Cep192 and Cenexin. Moreover, we uncover that replication-origin firing and S-phase progression specifically depends on the centrosomal activator Cep192, which initiates Plk1 activation both at centrosomes and the cytoplasm.

Our results indicate that the cytoplasmic and centrosomal pools of Plk1 activity are under distinct control. While in G2 Bora is the main activator of the cytoplasmic Plk1 pool, Cep192 and Cenexin are largely responsible for the activity of the centrosomal Plk1 pool. This indicates that both the Plk1 pools must be considered separately. This is in agreement with previous studies which indicated that DNA-damage recovery is under the control of cytoplasmic Plk1, but not nuclear Plk1, or a Plk1 version unable to diffuse from centrosomes^15^; or that kinetochore-microtubule attachments are regulated specifically by a kinetochore-bound form of Plk1^61^. Nevertheless, our FRET and FRAP measurements also indicate partial diffusion of Plk1 protein and activity between the cytoplasmic and centrosomal pools, consistent with previous work^62, 63^. Indeed, our data revealed an exchangeable pool and an immobile Plk1 tool at centrosomes that depends on Cep192 and Cenexin, in contrast with a previous study reporting only an exchangeable pool^63^. This difference may reflect different Plk1 expression levels as we studied the endogenous protein, instead of an exogenously tagged version used previously.

A second major conclusion of this study is that each Plk1 activator differentially regulates the Plk1-dependent cell-and centrosome cycle transitions. While Bora is the main driver of mitotic entry, DNA-damage recovery, and centrosome maturation, we uncover a major role of Cep192 in S-Phase progression and centriole disengagement, and a restricted role for Cenexin in centriole disengagement. Earlier studies reported negligible Plk1 activity during early cell cycle phases and proposed that Plk1 activation is restricted to G2, requiring S-phase completion^2, 17, 21^. Other studies, however, provided evidence for a role of Plk1 in DNA replication^5, 36–38^. Our data indicate that human cells exhibit low, but functionally significant Plk1 activity in S-Phase that drives DNA replication-origin firing. This S-phase activity depends on Cep192, which is thought to recruit Aurora A and Plk1 to centrosomes to drive centrosome maturation and spindle formation^26, 64^, but does not require centrosomes themselves. The most logic conclusion is that Plk1 activity in S-phase depends mainly on Cep192, but that this interaction is not dependent on this particular localization. Whether this S-phase activation depends on Aurora-A, remains to be determined.

As cells progress to G2, Bora becomes the main Plk1 activator, consistent with parallel Aurora A activation^65^. Our data align with previous findings on the critical role of Plk1 in DNA damage checkpoint recovery, centrosome maturation and mitotic entry in late G2^22, 25, 50^. We observed a similar magnitude of impaired centrosome maturation after depletion of either Bora or Plk1, implying that Cep192 and Cenexin cannot substitute for cytoplasmic Bora during this process, despite their centrosomal localization. We still observed a partial impairment of centrosome maturation after Cep192 depletion, consistent with its direct role in centrosome-cycle progression^66–69^, but this activity alone is insufficient to drive centrosome maturation. This suggest that full centrosome maturation, may require Plk1 to phosphorylate additional regulators beyond those at centrosomes.

Finally, Plk1 regulates centriole disengagement^52, 54^, a critical step for licensing centrioles for the next duplication round. This appears to be the only step that depends on the Cep192-and Cenexin-dependent centrosomal Plk1 pool. Our data indicates that Cep192 is the primary activator for centrosomal Plk1 and that Cenexin only has limited contribution towards fine-tuning centrosomal Plk1 activity. This aligns with a recent studies revealing how Cenexin fines-tunes centrosome maturation on old and new centrosomes through Plk1 activity^30, 70^. Centriole disengagement which normally occurs during telophase, thus appears to be predominantly regulated by Cep192- and Cenexin-dependent Plk1 pools, as Bora is degraded in mitosis^21, 71, 72^. This is also consistent with our previous finding that the Plk1 activity threshold required for centriole disengagement is much lower^55^ than that required for Plk1-dependent other mitotic steps.

Overall, we demonstrate that that Plk1 activity is regulated by Bora, Cep192 and Cenexin with each required for distinct cell cycle stages. This stage-specific regulation is reminiscent of the Cdk– cyclin system, where cyclin expression controls Cdk temporal activity and drive cell cycle progression, suggesting that Plk1, like Cdks, rely on a sequential activation mechanism to coordinate stage-specific functions. Similar to D-type cyclins, which are present through several cell cycle stages, Cep192 is present early and persists until mitotic exit, while Bora resembles more Cyclin-E, as it plays a crucial role during a particular cell transition, G2 and mitotic entry (see model in Fig. 8F). This differential regulatory network can, however, induce genetic instability, as observed with Bora depletion, where the mitotic progression is delayed while the centrosome cycle advances, causing a severe desynchrony between the two otherwise synchronous processes. It therefore will be exciting to explore in the future, whether such type of asynchronies can also be observed in pathological conditions, such as genetically instable cancer cells.

## Methods

### Cell culture and drug treatments

hTERT-RPE1 cells (ATCC: CRL-4000), hTERT-RPE1 Plk1-EGFP, hTERT-RPE1 USP28^-/-^ (kind gift by Arshad Desai, University of California, San Deigo, United States)^43^, h-TERT RPE1 Cenexin^-/-^ (kind gift by gift of Brian Tsou, Sloan Kettering Institute, United States)^73^, hTERT-RPE1 mRuby-PCNA (kind gift by Alexis Barr, Imperial College London, United Kingdom)^74^, hTERT-RPE1 Cyclin A2-mScarlet + Plk1-FRET Sensor, hTERT-RPE1 Cyclin A2-mScarlet + Plk1-FRET-PACT sensor, hTERT-RPE1 Separase Sensor cells were cultured in Dulbecco’s Modified Eagle’s Medium (Thermo-Fisher Scientific: 61965-026) supplemented with 10% foetal calf serum (FCS) [Regular: (Thermo-Fisher Scientific: A5256701) or Tet-System Approved: (Thermo-Fisher Scientific: A4736201) as required] and 100U/ml of each penicillin and streptomycin (P/S) (Thermo-Fisher Scientific: 15140122) at 37°C, 95% relative humidity and 5% CO_2_ in humidified CO_2_ incubator. All cell lines were routinely tested for mycoplasma contamination by PCR. For live-cell imaging, cells were cultured in Leibovitz’s L-15 medium without Phenol Red (Thermo-Fisher Scientific: 21083-027) with 10% FCS and 100 U/ml P/S at 37°C. Cells were treated with following inhibitors/drugs to inhibit/induce indicated proteins as per experimental requirement, Cdk1: 9 µM RO3306 (Sigma Aldrich: SML0569), Plk1: 10 nM BI2536 (Selleck Chemicals: S1109), 300 nM Plk4: Centrinone (Tocris Bioscience: 5687), Eg5: 10 µM (+)-S-Trityl-L-cysteine/STLC (Sigma Aldrich: 164739), DNA polymerase (to induce mild replication stress): 400 nM Aphidicolin (Sigma Aldrich: A0781) and 1 µg/ml Doxycycline (Sigma Aldrich: D3447-1G). To label microtubules and DNA in live cell imaging 50 nM SiR-Tubulin (Cytoskeleton Inc: CY-SC002), SiR-DNA (Cytoskeleton Inc: CY-SC007) and SPY505-DNA (Cytoskeleton Inc: CY-SC101) were used according to manufacturer’s instructions.

### Preparation of stable cell lines

hTERT-RPE1 Separase sensor cells were prepared by transfecting hTERT-RPE1 FRT/TR cells with 0.5 µg Separase Sensor (pcDNA5-FRT/TO-mCherry-Scc1^(142–467)-ΔNLS^-EGFP-PACT)^60^ and 4.5 µg pOG44 plasmids using X-tremeGENE™ 9 (Merck: XTG9-RO) transfection reagent according to manufacturer’s instructions. The transfected recombinants were selected with 200 µg/ml Hygromycin-B (Invivogen: ant-hm-5) in DMEM with 10% FCS and 100 U/ml P/S. A two-tiered approach was used to prepare hTERT-RPE1 Cyclin A2-mScarlet + Plk1-FRET Sensor and hTERT-RPE1 Cyclin A2-mScarlett + Plk1-FRET-PACT (Centrosome localised) sensor cells, where first hTERT-RPE1 FRT/TR cells were transfected with 0.5 µg Cyclin A2 (pcDNA5-FRT/TO-Cyclin A2-mScarlet) and 4.5 µg pOG44 plasmids as described above to generate hTERT-RPE1 Cyclin A2-mScarlet cells. These cells were later transfected again with either Plk1-FRET sensor c-jun substrate plasmid ^32^ (Addgene Plasmid # 45203) or c-jun based Plk1 FRET sensor tagged to PACT domain at c-terminus enabling centrosomal localisation ^33^ (Addgene Plasmid # Plasmid #106907) using X-tremeGENE™ 9 (Merck: XTG9-RO) transfection reagent according to manufacturer’s instructions. Transfected cells were selected with DMEM with 10% FCS (Tet-System compatible) and 100U/ml P/S supplemented with both 600 µg/ml of G418 (Invivogen: ant-gn-5) and 200 µg/ml Hygromycin-B (Invivogen: ant-hm-5) followed by single cell cloning. Centrosome devoid USP28^-/-^ cells were prepared by by treating the cells with 300nM of the Plk4 inhibitor centrinone ^42^ followed by single-cell sorting by FACS and immunofluorescence-based selection of clones devoid of centrioles (Fig. S4). These cells were constantly maintained in 300nM centrinone to prevent de-novo centriole formation. For generating Plk1-EGFP knock-In cell line, hTERT-RPE1 cells were co-transfected with Cas9 eGFP, sgRNA 5′ TCGGCCAGCAACCGTCTCA 3′ (targeting Plk1) and a repair template (Genewiz). The repair template was designed as a fusion of 5xGly eGFP flanked by two 500 bp arms, homologous to the genomic region around the Cas9 cutting site. 5 days after transfection EGFP positive cells were sorted and expanded for 1 week before a second sorting of single cells in a 96 well plates. After 2–3 weeks cells were screened by PCR.

### RNA Interference

siRNA transfections were performed using Lipofectamine RNAiMAX (Thermo-Fisher Scientific: 13778075) according to manufacturer’s instructions. RNAi was performed for 72 h; when combined with inhibitors the drugs were added 60 hours post transfection. All the siRNA sequences used in the study were previously validated sequences: siControl (Qiagen, GGACCTGGAGGTCTGCTGT), siBora (Dharmacon, TAACTAGTCCTTCGCCTATTT) ^71^, siCep192 (Dharmacon, AAGGAAGACATTTTCATCTCTTT), siCenexin (Dharmacon, GGCACAACATCGAGCGCAT), siCyclin A2 SMARTpool (Dharmacon, GGAAATGGAGGTTAAATGT, TAGCAGAGTTTGTGTACAT, ATGAGGATATTCACACATA, TGATAGATGCTGACCCATA), siAurora-A (Dharmacon, ATGCCCTGTCTTACTGTCA), siPlk1 (Dharmacon, CGAGCTGCTTAATGACGAG), siSeparase (Dharmacon, GCTTGTGATGCCATCCTGA) ^75^.

### Antibodies

The following antibodies were used in this study: Mouse anti-α-tubulin (Geneva antibody facility: AA345-M2a; 1:250: ExM) ^76^, Mouse anti-β-tubulin (Geneva antibody facility: AA344-M2a; 1:250: ExM) ^76^, Mouse monoclonal anti-α-tubulin (Clone: DM1α, Sigma Aldrich, T9026, 1:5000: Western Blotting), Rabbit polyclonal anti-Bora (gift from Erich Nigg, 1:1000 Western blotting), Rabbit polyclonal anti-CEP192 (Bethyl: A302-324A, 1:1000 Western blotting), Rabbit polyclonal anti-Cenexin (abcam: ab43840, 1:1000 Western blotting) , Mouse monoclonal anti-Aurora A (Clone 4/IAK1, BD Biosciences: 610939, 1:1000 Western blotting), Mouse monoclonal anti-Plk1 (Clone: 36-298, abcam: ab17057, 1:1000 Western blotting), Rabbit polyclonal anti-Pericentrin (abcam: ab4448; 1:250: ExM, 1:2000: IF), Mouse monoclonal anti-Cyclin A2 (Clone: E32.1, abcam: ab38; 1:1000: Western Blotting), Chicken polyclonal anti-GFP (Thermo-Fisher Scientific: A10262; 1:2000 IF), Mouse monoclonal anti-Centrin (Clone: 20H5, Merck Millipore: 04-1624 ; 1:1500 IF), Rabbit polyclonal anti-53BP1 (Novus Biologicals: NB100-304: 1:2000 IF), Mouse monoclonal anti-γ-tubulin (Clone: GTU-88, Sigma Aldrich: T6557, 1:2000 IF), Mouse monoclonal anti-PCNA (Clone: PC10, Santacruz Biotechnology, SC-56), Mouse monoclonal anti-γ-H2AX pSer139 (Clone: JBW301, Merck Millipore, 05-636) and Mouse monoclonal anti-Separase (abcam: ab16170; 1:500 Western blotting). All the Alexa Fluor-conjugated secondary antibodies were purchased from Thermo-Fisher Scientific and used at 1:500 dilution. For Immunoblotting experiments, Horseradish Peroxidase (HRP)-Conjugated goat anti-mouse antibody was purchased from Thermo-Fisher (Cat # 32430) and HRP-Conjugated goat anti-rabbit antibody was purchased from Bio-Rad (Cat# 1706515) and both of them were used at 1:10,000 dilution.

### FRET assay to measure Plk1 activity

Tetracycline inducible hTERT-RPE1 Cyclin A2-mScarlett Cells stably and constitutively expressing Plk1 FRET sensor were seeded in four-well glass bottom µ-Slide Ibidi chambers (Ibidi; 80426) and treated with different inhibitors/siRNAs as indicated in DMEM with 10% FCS and 1% P/S. The DMEM was replaced with Leibovitz L-15 supplemented with 10% FCS and 1% P/S containing same inhibitors as before if any. The chambers were acclimatised in 37 °C chamber before imaging. The acquisition was performed with an EC Plan Apochromat 100X (NA 1.56) oil objective on a Zeiss Cell Observer.Z1 spinning disk microscope (Nipkow Disk) equipped with a 37 °C chamber and a CSU X1 automatic Yokogawa spinning disk head. To perform FRET experiments samples were illuminated with 445 nm laser and the emission signal was split equally using DV2 split view system and CFP and YFP emissions were recorded on the split beams. 512×512-pixel size images were acquired with an Evolve EM512 camera (Photometrics) using Visiview 4.00.10 software. Acquired images were analysed using ImageJ to calculate the YFP to CFP emission intensity ratio after background subtraction.

### Immunofluorescence

Cells treated with indicated inhibitors/siRNAs were fixed in prechilled methanol at -20°C for 6 min. Fixed cells were blocked in Blocking solution (5% BSA in PBS), followed by incubating with indicated primary and secondary antibodies for 1 h and 30 min, respectively, with three washes of 10 min with PBS in between. Stained coverslips were mounted on glass slides with Vectashield with DAPI (Vector Laboratories: H-1200-10) and visualised using 100x PLAN Apochromat oil-immersion (NA 1.4) objective on an Olympus DeltaVision microscope (GE Healthcare) equipped with a DAPI/FITC/Rhodamine/CY5 filter set (Chroma Technology Corp) and a CoolSNAP HQ camera (Roper-Scientific). The three-dimensional image stacks were deconvolved with SoftWorx (GE Healthcare). The acquired images were cropped and processed with ImageJ (NIH) software to generate intensity projections (maximum/average) and extract fluorescence intensity and pixel size values according to the analysis requirement.

### Immunoblotting

Cells were grown in 60 mm plastic dishes and treated with inhibitors/drugs overnight. To prepare protein lysate the cells were scrapped off using cell scrapper and lysed in RIPA buffer (50 mM Tris pH-7.4, 150 mM NaCl 1% Nonidet P-40 (Thermo-Fisher Scientific: 85124), 0.5% Sodium deoxycholate (Sigma Aldrich: D5670), 0.1% Sodium dodecyl-sulphate in ultrapure water) supplemented with Protease inhibitor (Roche: 11873580001) and Phospho-STOP (Roche: 04906845001). Protein concentrations in the lysates were determined using the Bradford Protein Assay (Thermo-Fisher; 23200). Samples with equal amounts of protein were mixed with 4X Laemmli buffer and heated to 95 °C for 5 min. Proteins were separated on a 10% SDS-polyacrylamide gels and transferred onto a 0.45 µm pore size nitrocellulose membrane (Macherey-Nagel GMBH: 741280) by wet blotting. Membranes were blocked with 5% non-fat dry milk in TBS 0.1% Tween20 (TBS-T) for 30 min. After blocking, membranes were incubated with primary antibodies overnight at 4 °C in TBS-T 5% non-fat dry milk. Membranes were washed three times with TBS-T and incubated 1 h with the appropriate peroxidase-conjugated secondary antibody in TBS-T 5% non-fat dry milk. The membranes were washed thrice with TBS-T and the bands corresponding to protein of interest were detected by chemiluminescence using the Amersham ECL Prime Western Blotting Detection Kit (GE Healthcare; RPN2232) in a Fusion FX7 Spectra Multispectral Imaging system (Witec AG, Switzerland).

### Live cell imaging and analysis

For live cell imaging experiments hTERT-RPE1 cells endogenously tagged with mRuby-PCNA were plated in glass bottom Ibidi chambers (Ibidi GMBH: Cat # 81158) and normal DMEM medium was replaced with L15 Leibovitz’s medium supplemented with 10% FCS, 100 U/ml P/S, 50 nM SiR-Tubulin and 100 nM SPY505-DNA prior to imaging. The cells were treated with indicated inhibitors/siRNAs and imaged at 37 °C on a Nikon Ti microscope equipped with a 60x NA 1.3 oil objective, DAPI/FITC/Rhodamine/CY5 (Chroma, USA) filter set, Orca Flash 4.0 CMOS camera (Hamamatsu, Japan) and the NIS software. Cells were recorded every 10 min for 48 h with z-slices separated by 1 μm, and 100 ms exposure per z-slice at wavelengths of 488 (525), 561 nm (615 nm) and 647 (670) excitations (emission). The time-lapse movies were analysed manually for the duration of S and G2 phases based on PCNA localisation using Imaris software (Bitplane Inc).

### Flow cytometry-based cell cycle profile analysis

Asynchronously growing cells in log phase were collected by trypsinisation in a microcentrifuge tube, counted and fixed with prechilled 70% ethanol and incubated overnight at -20°C. Next day, the cells were washed with 1X PBS and resuspended in PI/RNase Staining Buffer (BD Biosciences: 550825) to a final concentration of 0.5 million cells per ml of suspension and incubated in dark for 1h at room temperature. The DNA content in the cells was analysed on CYTOFLEX flow cytometer (Beckman Coulter) using 561 nm laser after removing the doublets. The acquired data was analysed in FlowJo 10 Software (BD Biosciences) to extract the percentage of cells in different cell cycle stages based on DNA content.

### Expansion microscopy

Cells were grown on 12 mm circular glass coverslips (Thermo-Fisher Scientific) and treated with required inhibitors/drugs overnight. Next day the coverslips were treated with Acrylamide (AA)-Formaldehyde (FA) Solution [1.4% AA (Sigma Aldrich: A4058) and 2% FA (Sigma Aldrich: F8775) in PBS] for 5 h at 37°C to prevent protein crosslinking. Coverslips were next subjected to gelation by incubation for 1 hr at 37°C with monomer solution [19% Sodium Acrylate (Sigma Aldrich: 408220), 10% Acrylamide, 0.1% Bisacrylamide (Sigma Aldrich: M1533), 0.5% Tetramethyl ethylenediamine-TEMED (Thermo Fisher: 17919), 0.5% Ammonium Persulfate (Thermo Fisher: 17874) in PBS]. Post gelation, the coverslips were treated with denaturation solution [50 mM Tris (Sigma Aldrich: 99362), 200 mM Sodium dodecyl sulphate (Axon Lab AG: A2572.0500), 200 mM Sodium Chloride (Axon Lab AG: A3597.1000) in Nuclease free water, pH: 9.0] for 15 min on a rocker shaker at room temperature to detach the gels from coverslips. The gels were heated at 95°C for 90 min in denaturation solution followed by three 30 min washes with water. The gels were incubated with PBS for 15 min followed by 3 hrs incubations with primary and secondary antibodies followed each by three 10 min washes at 37°C and gentle shaking. Stained gels were kept overnight in water for optimal expansion. The size of gel was measured to calculate the expansion factor and the gel was cut into small pieces and placed in 2 well plastic bottom ibidi chamber (Ibidi GMBH: Cat # 80286)-The 3D image stacks of centrioles in G2 phase cells (4 centrioles in one cell) were acquired in 0.1 µm steps using a 100x PLAN-Apochromat oil-immersion (NA 1.4) objective on an Olympus DeltaVision microscope (GE Healthcare) equipped with a DAPI/FITC/Rhodamine/CY5 filter set (Chroma Technology Corp) and a CoolSNAP HQ camera (Roper-Scientific). The three-dimensional image stacks were deconvolved with SoftWorx (GE Healthcare). The acquired images were cropped and processed with ImageJ (NIH) software to construct 3D image to analyse the configuration of centrioles (orthogonal orientation and distance in between) for each image.

### Separase Activity Measurements

Separase activity measurements were performed on live cells. hTERT-RPE1 cells expressing Separase sensor were plated in glass bottom Ibidi chambers (Ibidi GMBH: Cat # 81158) and DMEM containing Tet-System approved FCS and transfected with indicated siRNAs. DMEM was replaced with L15 Leibovitz’s medium supplemented with 10% FCS, 100 U/ml P/S, 50 nM SiR-DNA and 1 µg/ml Doxycycline 3 hrs prior to imaging. Cells with two distinct centrosomes and intact nucleus (G2 phase) were selected and 3D image stacks of centrosomes were acquired for FITC, TRITC and Cy5 channels in 0.1 µm incremental z-steps using a 100x oil-immersion (NA 1.4) objective on an Olympus DeltaVision microscope (GE Healthcare) equipped with a DAPI/FITC/Rhodamine/CY5 filter set (Chroma Technology Corp) and a CoolSNAP HQ camera (Roper-Scientific). The three-dimensional image stacks were deconvolved with SoftWorx (GE Healthcare). The acquired images were cropped and processed with ImageJ (NIH) software to extract the intensities of GFP and mCherry in the centrosomes for each image and plotted as the ratio of GFP to mCherry as the measure of Separase activity.

### Statistical analysis

Statistical tests for all figures were performed using GraphPad Prism 10 (GraphPad), the statistical tests employed in every case are described in the figure legends. Minimum three independent biological replicates were performed in all experiments.

## Acknowledgements

Authors thank M. Gotta (University of Geneva, Switzerland) and A. Desai (University of California, San Diego, United States) for cell lines, E. Schiebel (University of Heidelberg) for plasmid constructs, P. Guichard, M. Laporte and V. Hamel (University of Geneva, Switzerland) for expansion microscopy support, Members of Bioimaging and FACS facility (University of Geneva, Switzerland) for experimental support, Monica Gotta and team (University of Geneva, Switzerland) and members of Meraldi laboratory for helpful discussions and support.

## Competing interests

The authors declare no competing or financial interests.

## Author contributions

Conceptualization: D.D., P.M.; Formal analysis: D.D., C.B., L.C. P.M.; Investigation: D.D., C.B. L.C. D.H.; Writing - original draft: D.D.; Writing - review & editing: D.D., P.M.; Visualization: D.D.; Supervision: P.M.; Project administration: P.M.; Funding acquisition: D.D., P.M.

## Funding

This work was supported by the Swiss National Science Foundation (Schweizerischer Nationalfonds zur FoCrderung der Wissenschaftlichen Forschung; SNF) project grant (No. 31003A_208052) to P.M., Novartis Foundation for Biomedical Research, Young Investigator Grant (No. 23A015) to D.D., Foundation NOVA Project Grant to D.D., and the Université de Genève.

## Figure Legends

**Figure S1:**
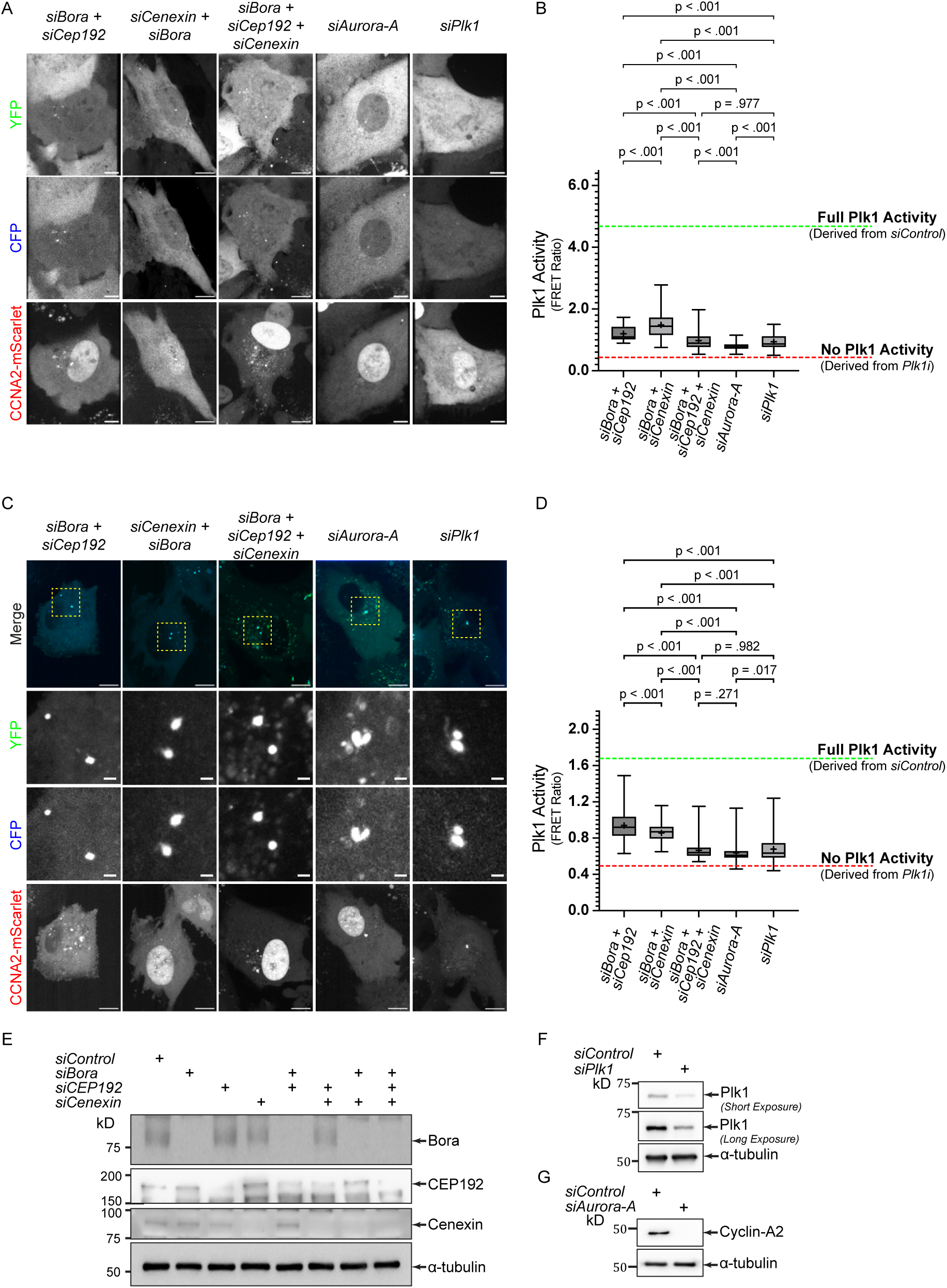
Different Plk1 activators activate distinct Plk1 pools. **(A)** Representative images of G2 phase hTERT-RPE1 cells expressing cellular Plk1 activity sensor and Cyclin A2-mScarlet treated with indicated siRNAs. **(B)** Box and Whisker plot for FRET measurement-based quantification of cellular Plk1 activity calculated from G2 phase cells represented in **(A)** derived from N=3 independent experiments comprising n= 120: siBora + siCep192, 135: siBora + siCenexin, 127: siBora + siCep192 + siCenexin, 124: siAurora and 121: siPlk1 cells. **(C)** Representative images of G2 phase hTERT-RPE1 cells expressing centrosomal Plk1 activity sensor and Cyclin A2-mScarlet treated with indicated siRNAs or inhibitors. **(D)** Box and Whisker plot for FRET measurement-based quantification of centrosomal Plk1 activity calculated from G2 phase cells represented in **(C)** derived from N=4 independent experiments comprising n= 181: siBora + siCep192, 187: siBora + siCenexin, 181: siBora + siCep192 + siCenexin, 182: siAurora and 182: siPlk1 cells. **(E-G)** Western Blot confirming the depletion of indicated proteins after indicated siRNA treatments. Data represent the entire range with values between first and third quartiles in boxes. ‘**+**’ sign in each column represent the mean value (p values for individual comparisons indicated on graphs: Kruskal-Wallis test). Scale bars = 5 µm.

**Figure S2:**
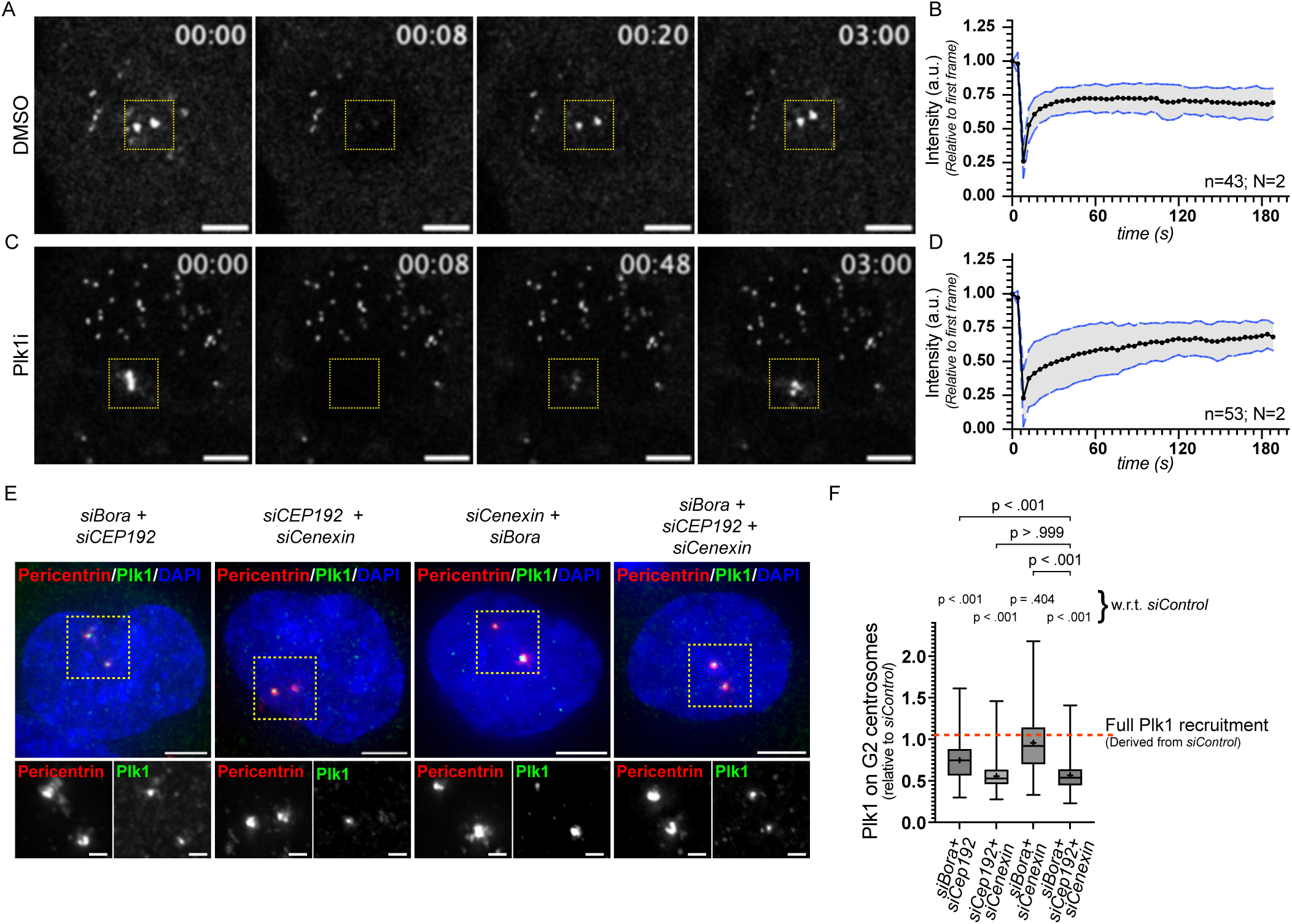
Plk1 activity does not affect the resident pool of Plk1 on centrosomes. **(A-D)** Representative time stamp images and quantifications from FRAP movies from G2 phase Plk1-EGFP tagged hTERT-RPE1 cells treated with DMSO **(A**: quantification of Plk1 recovery on centrosomes in **(B))** and Plk1i (**(C**: quantification of Plk1 recovery on centrosomes in **(D))**. Data obtained from N=3 independent experiments consisting of n=43 and 53 DMSO and Plk1i treated cells, respectively. **(E)** Representative immunofluorescence images of hTERT-RPE1: Plk1-EGFP cells treated with indicated siRNAs and stained for Pericentrin (Red) and EGFP (Green). **(F)** Box and Whisker plot for measurement of Plk1 levels on centrosomes during G2 phase under the conditions indicated in **(E)** generated from N=5 independent experiments comprising n= 276: siBora + siCep192, 287: siCep192 + siCenexin, 286: Bora + siCenexin and 282: siBora + siCep192 + siCenexin cells. Data presented as the entire range with values between first and third quartiles in boxes. ‘**+**’ sign in each column represent the mean value (p values for individual comparisons indicated on graphs: Kruskal-Wallis test). Scale bars Images: 2 µm and zoomed insets: 0.5 µm.

**Figure S3.**
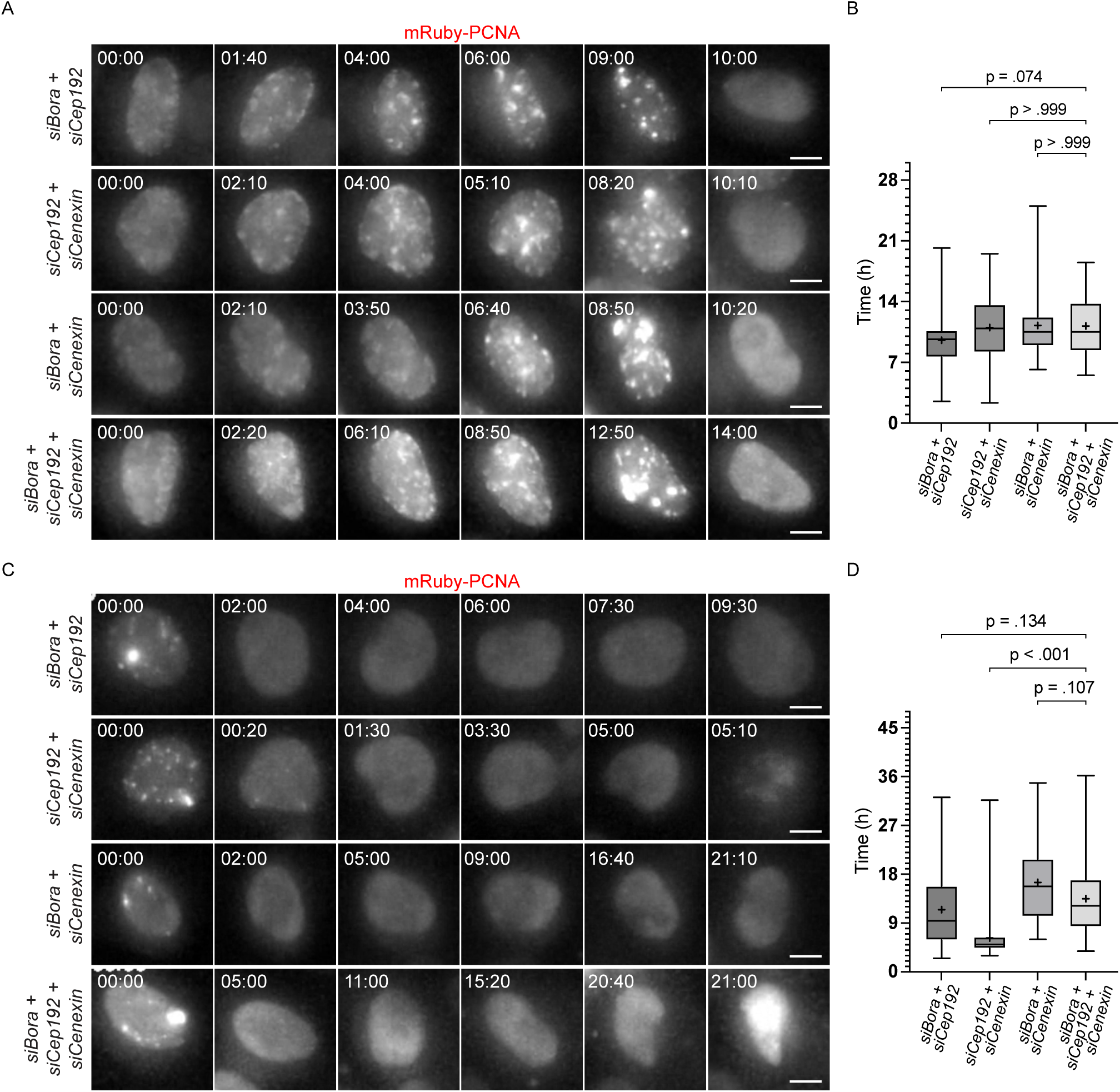
Perturbing Plk1 activity delays the progression through S and G2 phase. **(A)** Live-cell time-lapse images of hTERT-RPE1: mRuby-PCNA cells treated with indicated siRNA combinations as they progress through S phase. **(B)** Box and whisker plot for measurement of the duration of S phase in cells after depletion of proteins indicated in **(A)**. Data derived from N=3 independent experiments comprising n= 45: siBora + Cep192, 52: siCep192 + siCenexin, 57: siBora + siCenexin and 51: siBora + siCep192 + siCenexin cells. **(C)** Live-cell time-lapse images of hTERT-RPE1: mRuby-PCNA cells treated with indicated siRNA combinations as they progress through G2 phase. **(D)** Box and whisker plot for measurement of the duration of S phase in cells after depletion of proteins indicated in **(A)**. Data derived from N=3 independent experiments comprising n= 82: siBora + Cep192, 56: siCep192 + siCenexin, 64: siBora + siCenexin and 71: siBora + siCep192 + siCenexin cells. Data presented as the entire range with values between first and third quartiles in boxes. ‘**+**’ sign in each column represent the mean value (p values for individual comparisons indicated on graphs: Kruskal-Wallis test). Scale bars: 5 µm.

**Figure S4.**
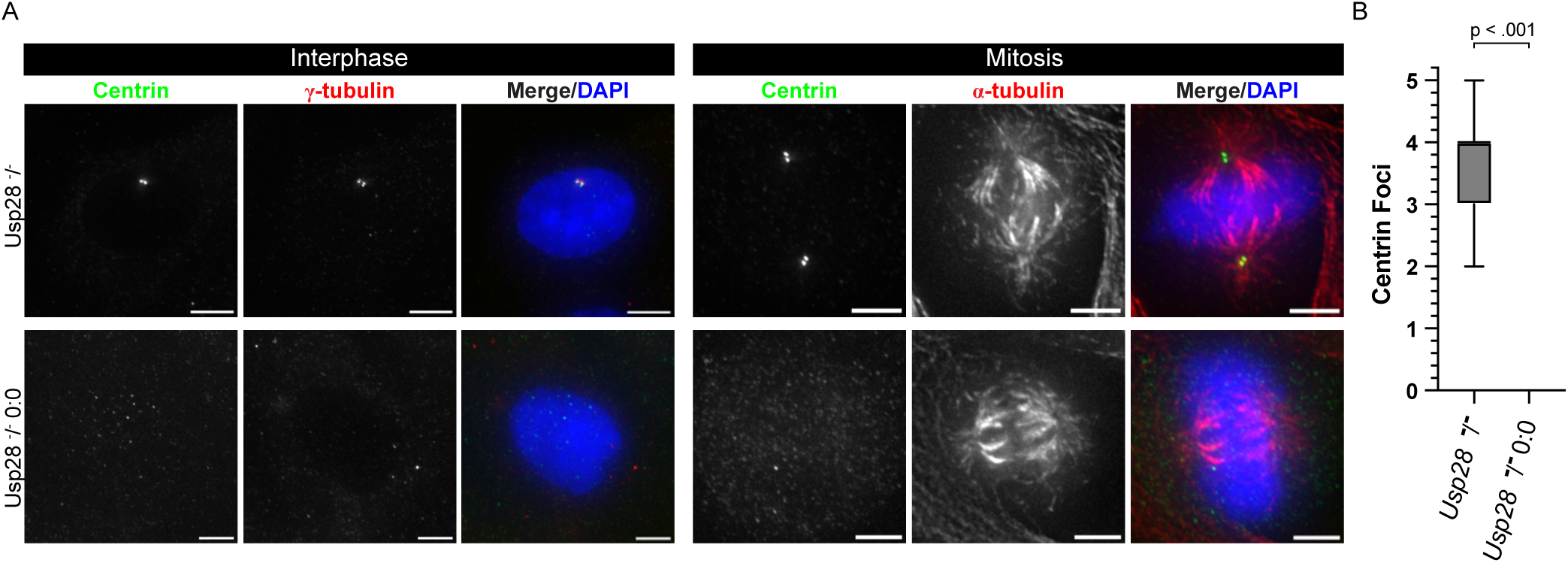
Centriole loss after prolonged centrinone treatment. **(A)** Representative images of interphase and metaphase hTERT-RPE1: Usp28 ^-^/^-^ and hTERT-RPE1: Usp28 ^-^/^-^ 0:0 cells stained for centrioles with Centrin (green) confirming absence of centrosomes. **(B)** Box and Whisker plot depicting the number of centrioles present in hTERT-RPE1: Usp28 ^-^/^-^ and hTERT-RPE1: Usp28 ^-^/^-^ 0:0 cells calculated from N=3 independent experiments with a total of n= 47 cells in each case. Data presented as the entire range with values between first and third quartiles in boxes. ‘**+**’ sign in each column represent the mean value (p values for the comparisons indicated on graph: Student’s t-test). Scale bars: 5 µm.

**Figure S5.**
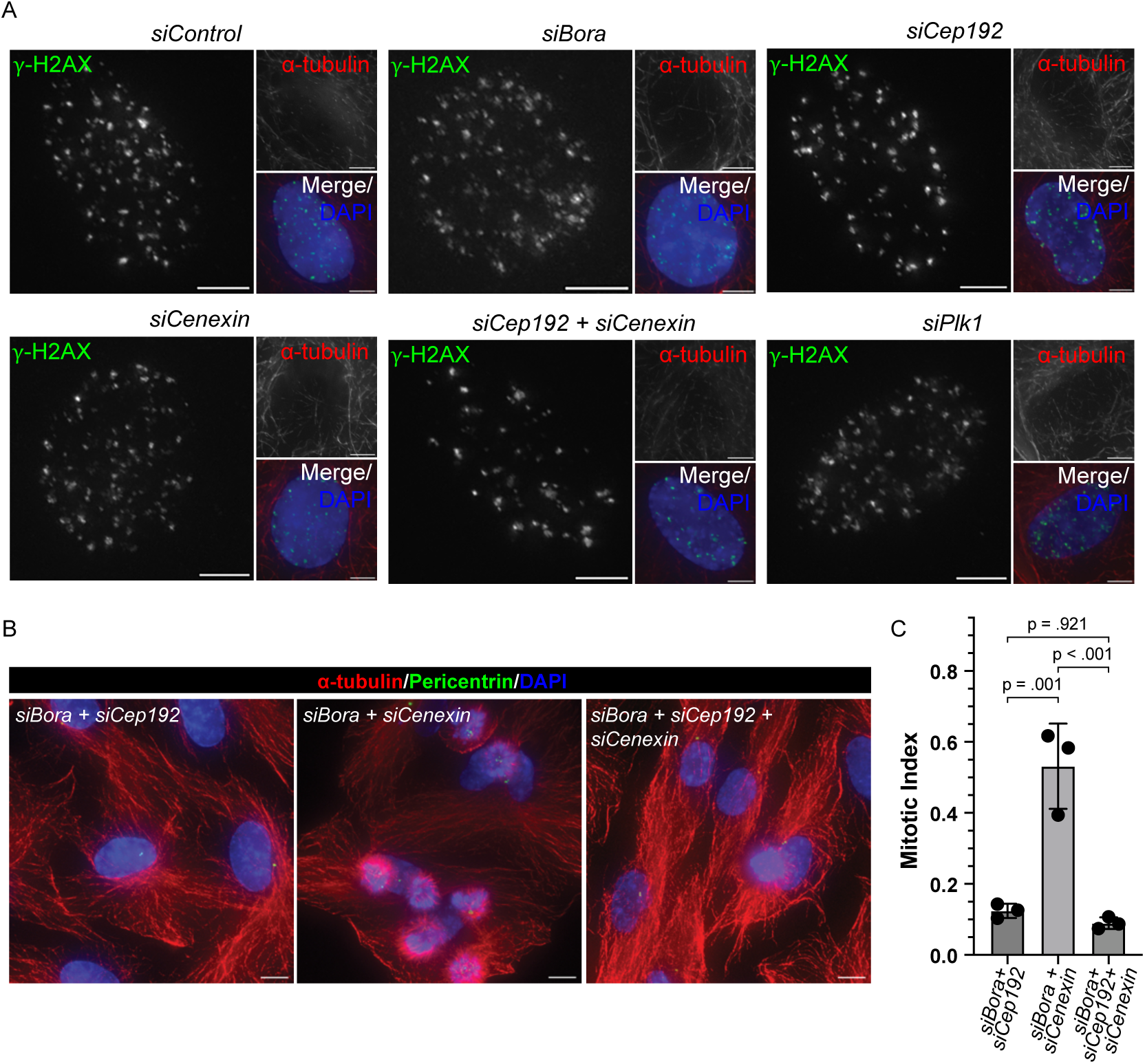
Confirmation of DNA damage induction by Doxorubicin pulse. **(A)** Representative images of synchronised and doxorubicin pulsed hTERT-RPE1 cells treated with indicated siRNA combinations and stained for γ-H2AX (green) to confirm induction of DNA damage after doxorubicin pulse. **(B)** Representative field images of hTERT-RPE1 cells treated with indicated siRNAs and stained for α-tubulin, Pericentrin and DAPI. **(C)** Quantification representing mitotic index in cells without doxorubicin pulse under indicated depletion conditions shown in **(B)** obtained from N=3 independent experiments consisting of n=591: siBora + siCep192, 955: siBora + siCenexin and 637: siBora + siCep192 + siCenexin cells. Data presented as the entire range with values between first and third quartiles in boxes. ‘**+**’ sign in each column represent the mean value (p values for individual comparisons indicated on graphs: Kruskal-Wallis test). Scale bars: 5 µm.

**Figure S6:**
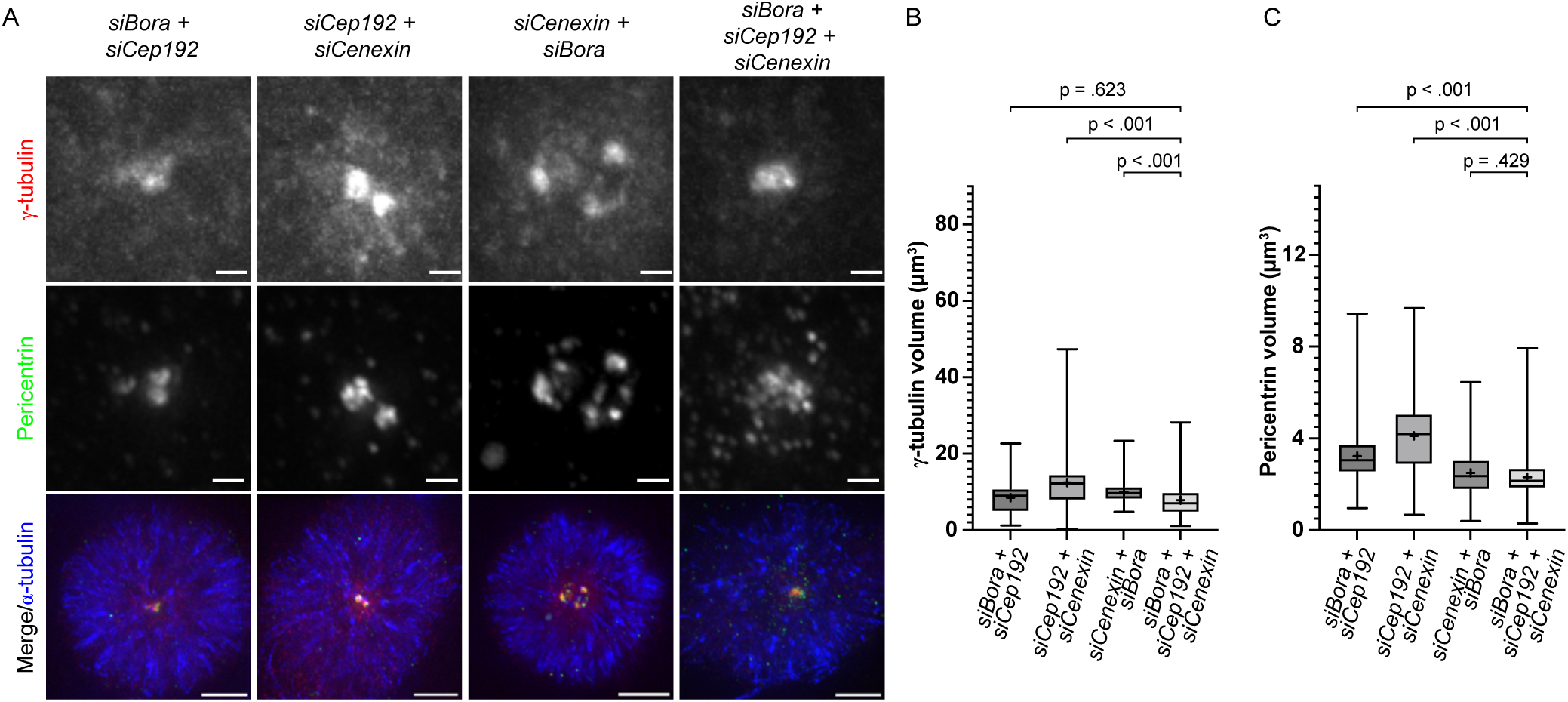
Plk1 activity is required for centriole maturation. **(A)** Representative images of G2 phase centrosomes stained with γ-tubulin (red) and Pericentrin (green) in hTERT-RPE1 cells treated with indicated siRNA combinations. **(B-C)** Box and Whisker plots representing the centrosomal levels of γ-tubulin **(B)** and Pericentrin **(C)** in cells represented in **(A)** as derived from N=3 independent experiments consisting of n= (138: siBora + siCep192, 164: siCep192 + siCenexin, 181: siCenexin + siBora and 167: siBora + siCep192 + siCenexin for γ-tubulin) and (n= 136: siBora + siCep192, 167: siCep192 + siCenexin, 181: siCenexin + siBora and 170: siBora + siCep192 + siCenexin for Pericentrin (green) cells. Data presented as the entire range with values between the first and third quartiles in boxes. ‘**+**’ sign in each column represent the mean value (p values for individual comparisons indicated on graphs: Kruskal-Wallis test). Scale bars: 0.5 µm.

**Figure S7.**
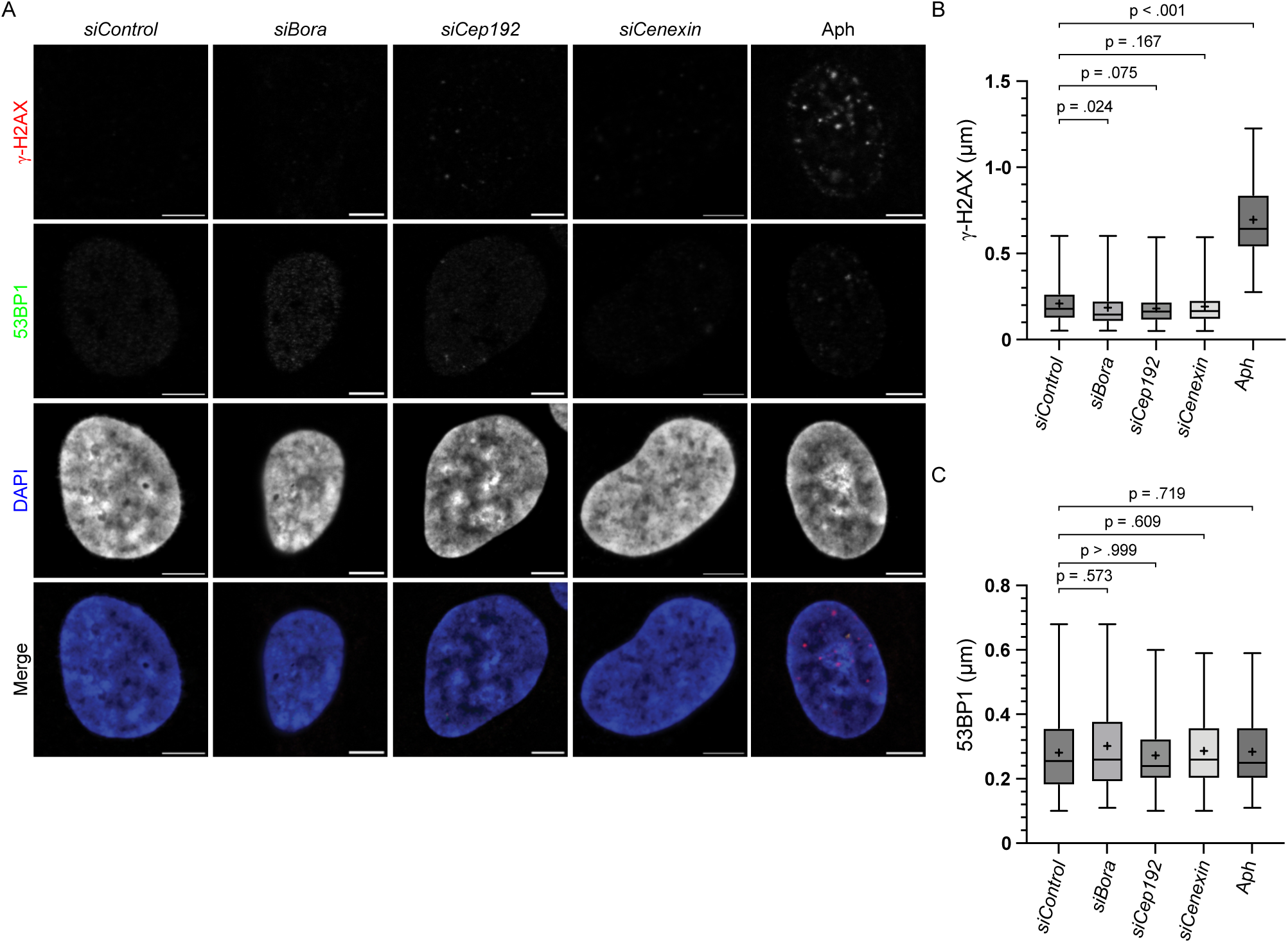
Depletion of Plk1 activators did not induce replication stress. **(A)** Representative images of hTERT-RPE1 cells treated with indicated siRNA/drugs and stained for indicated DNA damage marker proteins. **(B)** Box and Whisker plots depicting the size of γ-H2AX nuclear puncta shown in **(A)** from N=3 independent experiments with n= 186: siControl, 184: siBora, 186: siCep192, 187: siCenexin and 166: Aphidicolin treated cells. **(C)** Box and Whisker plots depicting the size of 53BP1 nuclear puncta shown in **(A)** from N=3 independent experiments with n= 228: siControl, 219: siBora, 169: siCep192, 223: siCenexin and 227: Aphidicolin treated cells. Data presented as the entire range with values between the first and third quartiles in boxes. ‘**+**’ sign in each column represent the mean value (p values for individual comparisons indicated on graphs: Kruskal-Wallis test). Scale bars: 0.5 µm.

## Supplementary Movies

**Movie 1:** FRAP recordings of hTERT-RPE1 expressing endogenously tagged Plk1-EGFP and treated with siControl. Every image was captured at an interval of 5s. The movie is shown at 3 frames per second (fps). Scale bars, 5 µm.

**Movie 2:** FRAP recordings of hTERT-RPE1 expressing endogenously tagged Plk1-EGFP and treated with siCep192. Every image was captured at an interval of 5s. The movie is shown at 3 frames per second (fps). Scale bars, 5 µm.

**Movie 3:** FRAP recordings of hTERT-RPE1 expressing endogenously tagged Plk1-EGFP and treated with siCenexin. Every image was captured at an interval of 5s. The movie is shown at 3 frames per second (fps). Scale bars, 5 µm.

**Movie 4:** FRAP recordings of hTERT-RPE1 expressing endogenously tagged Plk1-EGFP and co-treated with siCep192 and siCenexin. Every image was captured at an interval of 5s. The movie is shown at 3 frames per second (fps). Scale bars, 5 µm.

**Movie 5:** FRAP recordings of hTERT-RPE1 expressing endogenously tagged Plk1-EGFP and treated with DMSO/Control. Every image was captured at an interval of 4s. The movie is shown at 3 frames per second (fps). Scale bars, 5 µm.

**Movie 6:** FRAP recordings of hTERT-RPE1 expressing endogenously tagged Plk1-EGFP and treated with Plk1i(BI2536). Every image was captured at an interval of 5s. The movie is shown at 3 frames per second (fps). Scale bars, 5 µm.

**Movie 7:** Live-cell imaging of hTERT-RPE1 expressing endogenously tagged mRuby-PCNA and treated with siControl progressing through S phase. Every image was captured at an interval of 10m. The movie is shown at 5 frames per second (fps). Scale bars, 5 µm.

**Movie 8:** Live-cell imaging of hTERT-RPE1 expressing endogenously tagged mRuby-PCNA and treated with siControl progressing through G2 phase. Every image was captured at an interval of 10m. The movie is shown at 5 frames per second (fps). Scale bars, 5 µm.

**Movie 9:** Live-cell imaging of hTERT-RPE1 expressing endogenously tagged mRuby-PCNA and treated with siCyclin A2 progressing through S phase. Every image was captured at an interval of 10 m. The movie is shown at 5 frames per second (fps). Scale bars, 5 µm.

**Movie 10:** Live-cell imaging of hTERT-RPE1 expressing endogenously tagged mRuby-PCNA and treated with siCyclin A2 progressing through G2 phase. Every image was captured at an interval of 10 m. The movie is shown at 5 frames per second (fps). Scale bars, 5 µm.

**Movie 11:** Live-cell imaging of hTERT-RPE1 expressing endogenously tagged mRuby-PCNA and treated with siBora progressing through S phase. Every image was captured at an interval of 10 m. The movie is shown at 5 frames per second (fps). Scale bars, 5 µm.

**Movie 12:** Live-cell imaging of hTERT-RPE1 expressing endogenously tagged mRuby-PCNA and treated with siCep192 progressing through S phase. Every image was captured at an interval of 10m. The movie is shown at 5 frames per second (fps). Scale bars, 5 µm.

**Movie 13:** Live-cell imaging of hTERT-RPE1 expressing endogenously tagged mRuby-PCNA and treated with siCenexin progressing through S phase. Every image was captured at an interval of 10m. The movie is shown at 5 frames per second (fps). Scale bars, 5 µm.

**Movie 14:** Live-cell imaging of hTERT-RPE1 expressing endogenously tagged mRuby-PCNA and co-treated with siBora and Cep192 progressing through S phase. Every image was captured at an interval of 10 m. The movie is shown at 5 frames per second (fps). Scale bars, 5 µm.

**Movie 15:** Live-cell imaging of hTERT-RPE1 expressing endogenously tagged mRuby-PCNA and co-treated with siCep192 and siCenexin progressing through S phase. Every image was captured at an interval of 10 m. The movie is shown at 5 frames per second (fps). Scale bars, 5 µm.

**Movie 16:** Live-cell imaging of hTERT-RPE1 expressing endogenously tagged mRuby-PCNA and co-treated with siBora and siCenexin progressing through S phase. Every image was captured at an interval of 10 m. The movie is shown at 5 frames per second (fps). Scale bars, 5 µm.

**Movie 17:** Live-cell imaging of hTERT-RPE1 expressing endogenously tagged mRuby-PCNA and co-treated with siBora, siCep192 and siCenexin progressing through S phase. Every image was captured at an interval of 10 m. The movie is shown at 5 frames per second (fps). Scale bars, 5 µm.

**Movie 18:** Live-cell imaging of hTERT-RPE1 expressing endogenously tagged mRuby-PCNA and treated with siPlk1 progressing through S phase. Every image was captured at an interval of 10 m. The movie is shown at 5 frames per second (fps). Scale bars, 5 µm.

**Movie 19:** Live-cell imaging of hTERT-RPE1 expressing endogenously tagged mRuby-PCNA and treated with siBora progressing through G2 phase. Every image was captured at an interval of 10 m. The movie is shown at 5 frames per second (fps). Scale bars, 5 µm.

**Movie 20:** Live-cell imaging of hTERT-RPE1 expressing endogenously tagged mRuby-PCNA and treated with siCep192 progressing through G2 phase. Every image was captured at an interval of 10m. The movie is shown at 5 frames per second (fps). Scale bars, 5 µm.

**Movie 21:** Live-cell imaging of hTERT-RPE1 expressing endogenously tagged mRuby-PCNA and treated with siCenexin progressing through G2 phase. Every image was captured at an interval of 10 m. The movie is shown at 5 frames per second (fps). Scale bars, 5 µm.

**Movie 22:** Live-cell imaging of hTERT-RPE1 expressing endogenously tagged mRuby-PCNA and co-treated with siBora and Cep192 progressing through G2 phase. Every image was captured at an interval of 10 m. The movie is shown at 5 frames per second (fps). Scale bars, 5 µm.

**Movie 23:** Live-cell imaging of hTERT-RPE1 expressing endogenously tagged mRuby-PCNA and co-treated with siCep192 and siCenexin progressing through G2 phase. Every image was captured at an interval of 10 m. The movie is shown at 5 frames per second (fps). Scale bars, 5 µm.

**Movie 24:** Live-cell imaging of hTERT-RPE1 expressing endogenously tagged mRuby-PCNA and co-treated with siBora and siCenexin progressing through G2 phase. Every image was captured at an interval of 10 m. The movie is shown at 5 frames per second (fps). Scale bars, 5 µm.

**Movie 25:** Live-cell imaging of hTERT-RPE1 expressing endogenously tagged mRuby-PCNA and co-treated with siBora, siCep192 and siCenexin progressing through G2 phase. Every image was captured at an interval of 10 m. The movie is shown at 5 frames per second (fps). Scale bars, 5 µm.

**Movie 26:** Live-cell imaging of hTERT-RPE1 expressing endogenously tagged mRuby-PCNA and treated with siPlk1 progressing through G2 phase. Every image was captured at an interval of 10 m. The movie is shown at 5 frames per second (fps). Scale bars, 5 µm.

## Notes

### Competing Interest Statement

The authors have declared no competing interest.

### Summary of Updates

We made some text changes to the manuscript and updated the ladt figure with our proposed model

